# Site Specific Hydration Contributes to the Thermal Stability of Proteins and Protein-Complexes

**DOI:** 10.64898/2026.10.26.761984

**Authors:** Robert L. Rider, Zhenyu Xi, Adam Hillaire, Joanna Powell, Amy Spelbring, Kacie Evans, Carter Lantz, Elena Scott, David Barondeau, Shannon Raab, David Clemmer, Arthur Laganowsky, David H. Russell

## Abstract

Water is integral to protein structure and stability, yet ensemble measurements rarely identify the hydration interactions associated with stabilization. Here, variable-temperature electrospray ionization ion mobility mass spectrometry, conformation-resolved thermodynamic analysis, and molecular dynamics simulations are combined to examine solvent-dependent thermal stability of chymotrypsin inhibitor 2 (CI-2), a model two-state folder, and the barnase-barstar (Bn-B*) complex, which contains a highly polar binding interface. Replacing H_2_O with D_2_O raises the thermal transition temperature of both systems by approximately 3 °C, favors compact conformations, and reduces the relative thermodynamic accessibility of extended states. The conformationally resolved enthalpy-entropy compensation reveals D_2_O increases enthalpy for CI-2 but reduces enthalpy for Bn-B* as a result of opposing effects on the individual subunits of the complex. Molecular dynamics simulations reveal that lower solvent-accessible surface areas and fewer protein-water hydrogen bonds in D_2_O coexist with longer water residence times at specific structural sites. In CI-2, the strongest changes localize to the second β-strand/C-terminal region; in Bn-B*, persistent hydration is concentrated at the charged protein-protein interface. Together, these results support a model in which D_2_O dependent stabilization is associated not only with the extent of hydration, but the site and lifetime of water interactions. More broadly, this work provides an experimental framework for resolving solvent-dependent conformational thermodynamics of protein unfolding and interactions, enabling direct comparison of free-energy landscapes across solution environments.

**TOC:** 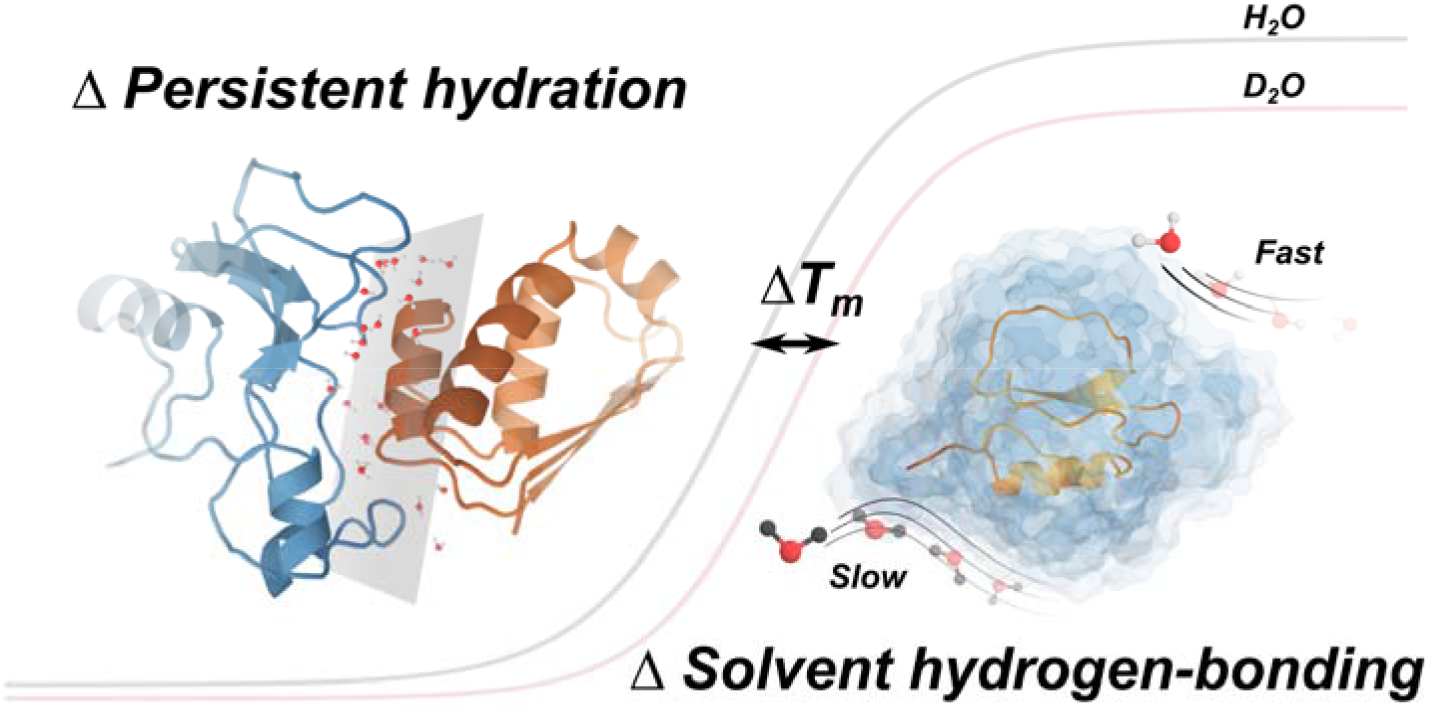

## Introduction

Protein stability reflects the balance between intramolecular interactions and interactions with the surrounding solvent.^1^ Although the role of the protein itself in determining conformational stability has been extensively studied, solvent interactions can also contribute substantially to the free energy of folding and unfolding. ^2^ ^3, 4^ Indeed, relatively small changes in solvent properties can shift protein conformational equilibria, highlighting the importance of the protein-solvent interface in determining stability.^5, 6^ Deuterium oxide (D_2_O) provides a useful perturbation of this balance, as proteins generally exhibit increased thermal stability in D_2_O relative to H_2_O.^7, 8^ While this effect has been attributed to differences in hydrogen bonding, solvent dynamics, and other physicochemical properties of the two solvents, the contribution of altered protein hydration remains less well defined.^9, 10^ ^11^

Chymotrypsin inhibitor 2 (CI-2) provides an established model for examining these effects. Classical biochemical and calorimetric studies characterized CI-2 as a highly cooperative, two-state folder, establishing a well-defined thermodynamic framework for its unfolding.^12^ ^13^ More recently, variable-temperature electrospray ionization ion mobility-mass spectrometry (vT-ESI-IM-MS) studies demonstrated that this apparent two-state transition comprises numerous discrete solution conformations with distinct temperature-dependent populations and thermodynamic properties.^4, 14^ These findings establish that the CI-2 unfolding landscape contains substantially greater conformational complexity than is apparent from ensemble measurements and provide an experimental framework for resolving how solvent perturbations influence individual conformers.

A complementary test is provided by barnase-barstar (Bn-B*), one of the best characterized protein-protein complexes. Its picomolar affinity is dominated by a highly polar, charge-complementary interface.^15, 16^ Structural and computational work has shown that interfacial water is integral to this surface. A single barstar D40A substitution is believed to perturb the interfacial water network which contributes to weakening of the complex binding by 7.7 kcal mol ¹.^17–22^ Simulations indicate strong interfacial bridging water that provides complex stability, with other very strong biomolecular interactions such as streptavidin-biotin also are reported to be influenced by hydration.^23, 24^ Bn-B* thus allows for comparison of the known hydration effects observed for protein unfolding to an intermolecular interface.

Here, we examine how solvent-dependent protein hydration alters the CI-2 unfolding and Bn-B* dissociation landscape by comparing temperature-dependent conformational populations in H_2_O and D_2_O. vT-ESI-IM-MS is combined with conformation-specific thermodynamic analysis, collision-induced unfolding, and molecular dynamics simulations to distinguish solvent-mediated effects from protein isotope effects. D_2_O increases the thermal stability, promotes compact conformations and decreases the thermodynamic favorability of extended states. Interestingly the thermodynamic signature shift (ΔH & -TΔS) is not consistent across both protein systems supporting a non-uniform explanation for the altered thermal stability. These changes coincide with reduced solvent exposure (fewer protein-solvent hydrogen bonds) in D_2_O as well as site specific increases in water-residence times. We conclude that the location and persistence of hydration, rather than hydration number alone, is an underappreciated contributor to protein thermal stability.

## Methods

### Materials

Recombinant chymotrypsin inhibitor 2 (CI-2) and barnase-barstar (Bn-B*) were expressed and purified in-house as described previously.^14, 22^ LC-MS grade water, ammonium acetate (AmAc), glacial acetic acid and deuterium oxide (D_2_O, 99.9% D) were purchased from Millipore Sigma (Burlington, MA). Nano-electrospray emitters were made using borosilicate glass and a P-1000 filament micropipette puller (Stutter Instruments, Novato CA).

### Sample Preparation

15 µM CI-2 solutions were buffer-exchanged into either D_2_O or H_2_O using Bio-Spin 6kDa MWCO SEC desalting columns (BioRad, Hercules CA) and left at 37□C for 24 h to allow for hydrogen deuterium exchange (HDX) to reach equilibrium (∼80%). Samples left for 30 days at 37□C saw no significant increase in HDX (Supporting Figure S1). Denaturing the protein to allow for ∼99% HDX and then refolding is likely to perturb the native conformations accessible and therefore, outside of the scope of our work. CI-2 samples were then diluted to 10 µM with a final concentration of 1% acetic acid (∼pH 2.6) for all experiments. 20 µM Bn-B* solutions were buffer-exchanged into 200mM AmAc in either D_2_O (pD 7)^25^ or H_2_O (pH 7) using Bio-Spin 6kDa MWCO SEC desalting columns and left at 37□C for 24 h to allow for HDX to reach equilibrium. Complex samples were diluted to a working concentration of 15 µM for all experiments.

### Variable-Temperature Ion Mobility Mass Spectrometry

Native protein conformational ensembles were interrogated using variable-temperature electrospray ionization (vT-ESI) coupled with traveling wave ion mobility mass spectrometry (IM-MS) on a Waters Synapt G2 from 5-90□C in 5□C increments for CI-2 and 25 - 90□C in 2.5□C increments starting at 55□C for Bn-B* (**Figure 1.1 & 1.2**). The vT-ESI source was modified slightly from the initial design of McCabe *et al*., to operate with the Waters MS.^26^ Supporting Figure S2 shows the calibration of the solution temperature inside the nESI emitter to ensure temperature readings were within ± 2□C. IM-MS settings were optimized to minimize collisional activation and preserve solution-like conformers in the gas phase as described previously.^27^ Parameters are listed in supporting Table S1.^27^ Drift times were converted to collision cross sections (CCS) using the denatured protein calibration protocol from Ruotolo *et al*.^28^

**Figure 1:**
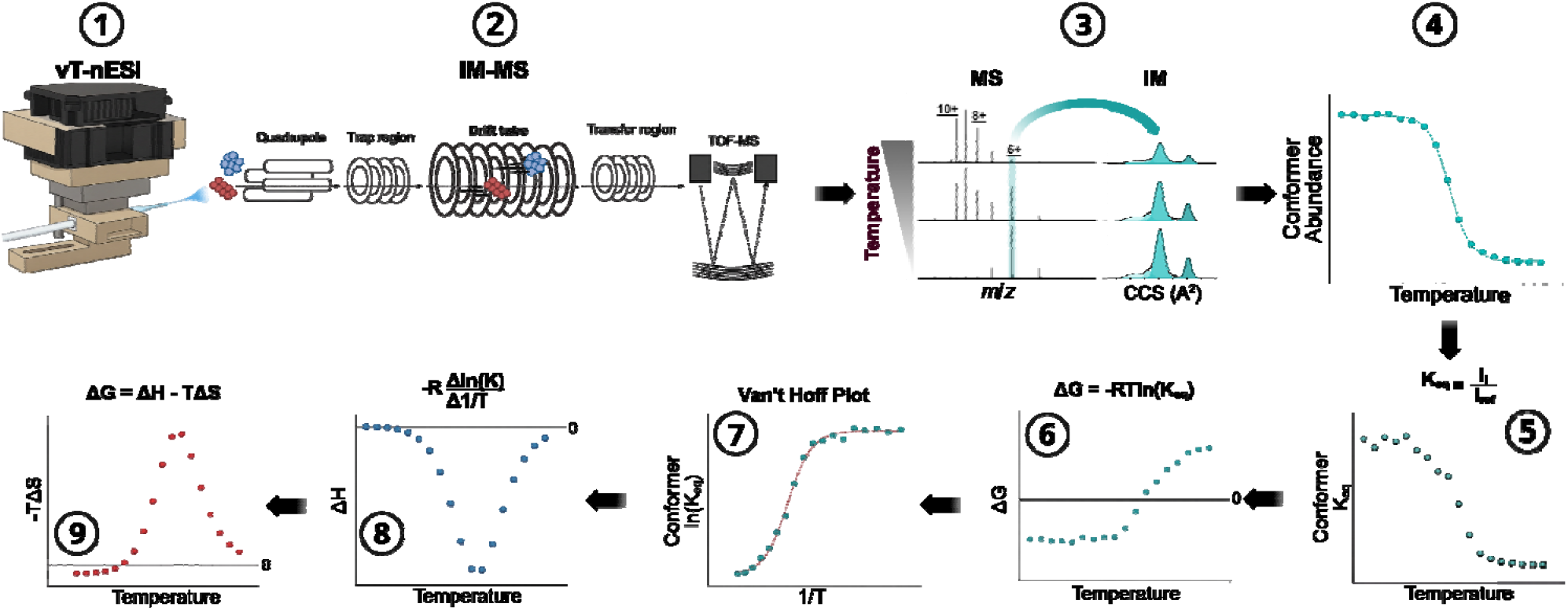
Experimental and analysis workflow for extraction of thermodynamic information (ΔG, ΔH, -TΔS) for eac conformation and temperature measured. Data and analysis for a dominant native conformation is shown. Image designed in BioRender.

### Data Processing and Conformation Assignment

Average charge state (Z_avg_) was calculated from the maximum intensity of each charge state in the CI-2 mass spectra using equation 1 where *z* is the charge state and *i* is the area intensity of the respective charge states. For each charge state (5+ through 10+), ion mobility CCS distributions were fit with the minimum number of Gaussians using a custom python script. An example of the fitting of the conformations within each charge state in both H_2_O and D_2_O solvents can be found in supporting figures S3 and S4. Weighted averages and standard deviations for CCS, and FWHM of each CI-2 conformation can be found in supporting Table S2. Population abundances of conformations were normalized across the entire distribution within each mass spectra collected to enable construction of temperature-dependent population profiles (**Figure 1.3 & 1.4**).

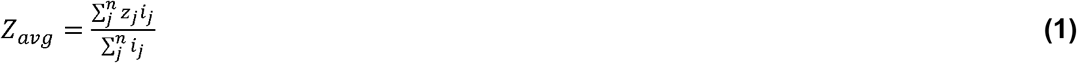

### Thermodynamic Analysis Using Van’t Hoff Relation

To extract thermodynamic parameters for individual features, we adapted and extended the analysis framework described previously for variable-temperature IM-MS thermodynamics.^4, 14^ Equilibrium constants were calculated using the smaller CCS conformation in the 7+ charge state as the reference state for CI-2 in equation 2 (**Figure 1.5**). This conformer was chosen since the 7+ charge is an intermediate state that only becomes present at experimental conditions (pH 2.6) and a lower abundance conformation present at all temperatures was found to be pertinent in the calculation of meaningful data. Selection of other conformations in the 6+, 7+, 8+ & 9+ charge states results in no change in ΔG sign (favorability) across temperature for most conformations or other behaviors that do not match the observed and expected trends seen with Z_avg_ changes and protein unfolding (see supporting figure S5). Selection of the reference conformation was done through an equally intensive process where the smallest conformation of the 5+ charge state of Bn was selected due to its relatively constant abundance across temperatures. It is important to note that the use of a reference conformation sets all thermodynamic parameters (ΔG, ΔH, -TΔS) for that conformer as 0, which is not biophysically true but is necessary to get relative thermodynamic values for all other conformations to that state.

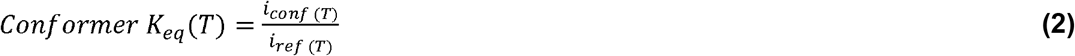

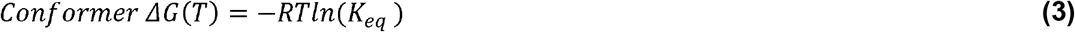

The Gibbs free energy (ΔG) at each temperature was calculated using the equilibrium constant and equation 3 (**Figure 1.6**). Unlike the previous studies that apply Gibbs Helmholtz analysis at the melting temperature and extrapolate to different temperatures, we implement a Van’t Hoff based analysis across the full temperature range.^29^ These different methods agree well at the inflection point as seen in supporting figure S6. The natural log of the equilibrium constant was plotted versus inverse temperature and fit with a sigmoidal distribution, which best represents the data (**Figure 1.7**). Using the derivative of the fit to the Van’t Hoff plot, enthalpy was calculated at each temperature using the known relation in equation 4 (**Figure 1.8**). Finally, the Gibbs equation (equation 5) was used to calculate entropy from the previously known ΔG and ΔH (**Figure 1.8**). Across all the temperatures measured, the conformations CCS and FWHM do not significantly change (see supporting Table S2), therefore it is reasonable to conclude that the heat capacity (measure for change in structure) for each conformation is 0. This is not to be misconstrued, as the protein system’s heat capacity does change during unfolding, however this analysis is focused on individual conformations and not the ensemble that is measured in traditional techniques such as CD or DSC, where a single heat capacity change is reported for the entire protein system.^30, 31^

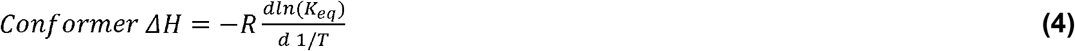

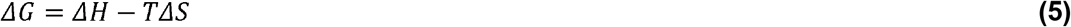

### Collision-Induced Unfolding Experiments

To distinguish solvent-driven stabilization from intrinsic isotope effects, collision-induced unfolding (CIU) experiments were performed on hydrogenated and perdeuterated forms of CI-2 and Bn-B*.^7^ Ions were subjected to collisional activation in the trap cell from 2-30V in 1V increments for CI-2 and 5-70V for the complex. Each mobility profile was fit using a custom python script to determine the abundance of each conformation and calculate the weighted average CCS. The change in average CCS was used to create a % unfolded curve versus activation voltage. Fitting the CIU curve using a sigmoidal equation, the midpoint was extracted and used as the CIU_50_ measurement.^32^

### Molecular Dynamics Simulations

To determine important regions and interactions for CI-2’s stability, molecular dynamic simulations (MDS) utilizing GROMACS^33^ with the CHARMM36^34^ force field and TiP3P water model were employed. Starting structures were derived from the crystal structure (PDB: 2CI2), with unresolved residues on the termini defined as loop regions. The structures were defined as 0 charge (pI= 6.43), placed in an 8nm box with water and allowed to equilibrate. All MD simulations were run at 300 K for 200 ns to allow sufficient sampling. Analysis was completed on the entire trajectory to calculate whole protein and residue specific information on solvent accessible surface area (SASA), radial distribution function with solvent and solvent-protein h-bond interactions. All MDS were completed for D_2_O as well using TiP3P heavy water model with no HDX taken into consideration to focus on the impact of solvent. Gas phase structures with their respective charge states (5+ - 10+) to match IM-MS data were calculated using a similar procedure as published prior.^35^ Briefly, protonation site was determined based on basicity and SASA, with the simulation being ran for 2000 ns at 363 K. CCS of the structures were calculated using IMPACT^36^ to best match experimental IM-MS data solution and gas phase MDS. Structures were visualized and processed in PYMOL.

For Bn-B* starting structures were derived from the PDB: 1BRS and ran at 300 K for 200 ns in both TiP3P light and heavy water after equilibration. The global SASA and hydrogen-bond distributions used the full 200 ns trajectories. Direct interface residues were defined by at least one interchain Cα pair within 8 Å in at least 10% of frames. For comparison with the published Bn-B* hydration analysis, water oxygen atoms within 3.5 Å of protein heavy atoms were classified as single-contact, multiple-contact, or bridging waters, and intermittent contact-correlation functions were computed.^17^ Chemically explicit one- and two-water bridges additionally required a donor-acceptor distance ≤3.5 Å and donor-H/D-acceptor angle ≥150°. H_2_O/D_2_O differences in global hydrogen-bond counts were evaluated framewise. PCA analysis was completed to determine the residues that contribute the most to the protein complex stability, which align very well with prior mutagenesis studies.^19^

## Results

Ensemble techniques such as isothermal titration calorimetry and circular dichroism have reported differences in protein behavior in deuterium oxide (D_2_O). Although D₂O is known to alter protein stability and dynamics, disentangling solvent, isotope, and intrinsic protein effects remains challenging.^7^ Recently, our group has used native mass spectrometry (nMS) to provide high-resolution insight into how altered hydration (H_2_O vs D_2_O) affects protein-ligand binding and conformational equilibria to various native and non-native states.^37–39^ Here, we first appl nMS to obtain conformation-specific thermodynamics of unfolding for CI-2, with the ability to resolve native, intermediate, and unfolded states within a single experiment. The same workflow is applied to the complex dissociation of Bn-B*, with the ability to discern contributions and hydration effects of the individual proteins. These results demonstrate the utility of nMS for probing hydration-dependent changes in protein physicochemical properties and conformational behavior.

### Solvent Dependent Unfolding and Dissociation

Temperature-dependent mass spectra reveal clear two-state unfolding behavior for CI-2 in both solvents, with systematic differences in charge-state distributions between H_2_O and D_2_O (**Figure 2A & 2B**). CI-2 in D_2_O consistently populates lower average charge states across the full temperature range (5-90 °C). At 5□C, H_2_O has a higher abundance of the 7+ charge state, and at higher temperatures >55□C, the 9+ is the dominant charge state in H_2_O, while the 8+ is the dominant charge state in D_2_O (Figure S7). These charge state differences indicate reduced solvent-accessible surface area (SASA) relative to H_2_O.^40, 41^ Average charge state (Z_avg_) analysis shows a reproducible increase in the apparent melting temperature in D_2_O by approximately 3 °C, consistent with enhanced thermal stability (**Figure 2C**). Importantly, intact hydrogen-deuterium exchange of ∼77% is observed, confirming solvent isotope incorporation of the most SASA regions (Figure S1). As CI-2 unfolds, the mass of the D_2_O sample increases, synonymous with an increase in SASA, supporting CI-2 being folded/native during the start of the experiments (Figure S8). Intentional denaturing to increase exchange closer to 100% were not completed as denaturing would likely perturb the native states accessible upon refolding and is beyond the scope of this work. The mass spectral data suggests that thermal stability of CI-2 is altered in D_2_O and could arise from the change in isotope exchange or solvent environment.

**Figure 2:**
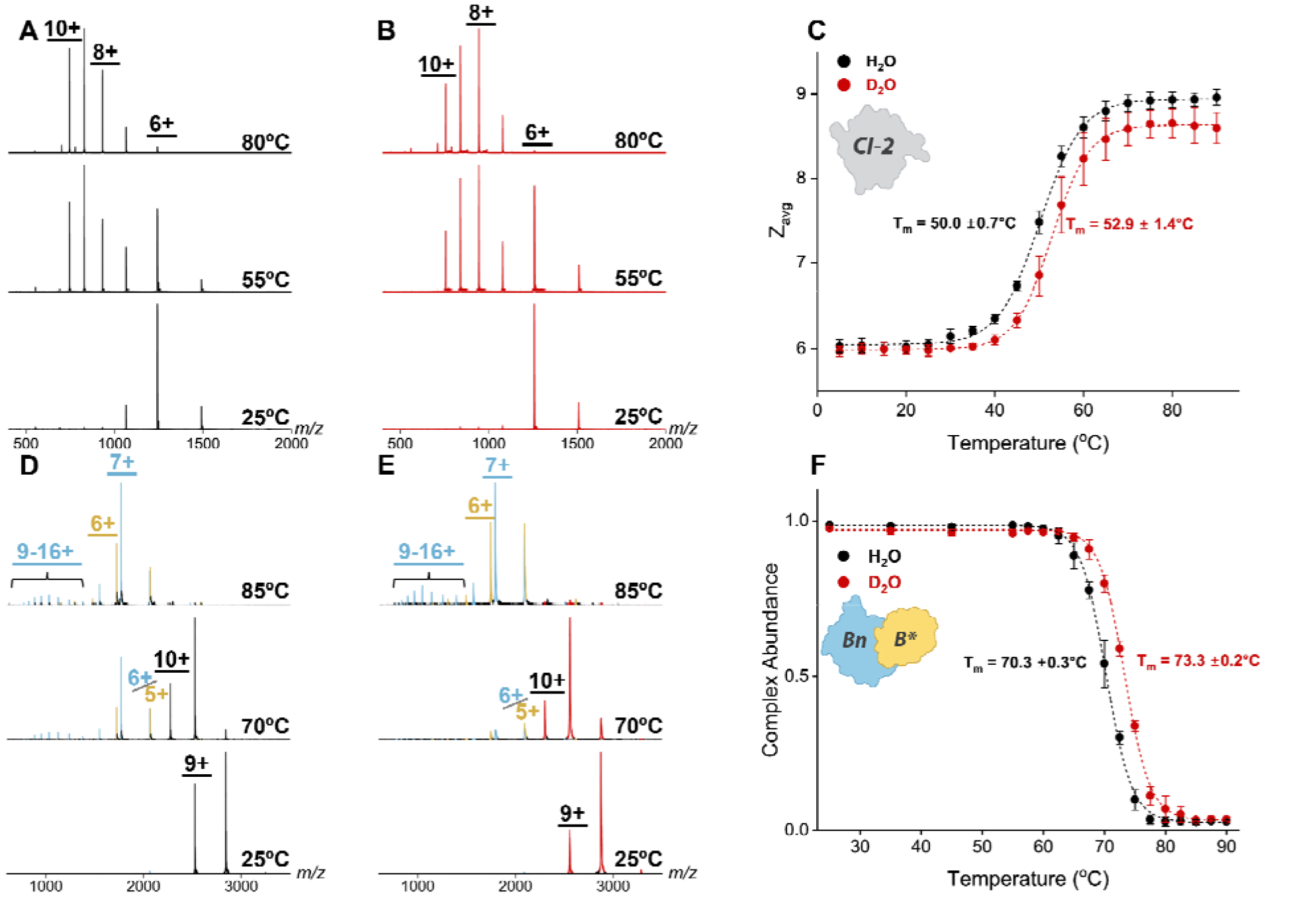
Representative mass spectra over the temperature range examined for CI-2 in A) H_2_O and B) D_2_O solvents. C) Average charge state analysis from 5-90□C is used to track the 2-state unfolding behavior of CI-2 in both solvents with the melting temperature being increased by ∼ 3□C in D_2_O (p-value < 0.05). Representative mas spectra over the temperature range examined for the Bn-B* complex in D) H_2_O and E) D_2_O solvents. Signals seen in blue represent Bn, yellow for B* and the complex is either black (H_2_O) or red (D_2_O) depending on the solvent system. D) Calculated complex abundance from 25-90□C used to track the complex thermal stability. The melting temperature is increased by 3□C in D_2_O (p-value < 0.01). Across all temperature ranges and protein systems D_2_O prefers a lower average charge state, indicating there is a difference in protein SASA between solvents. Error bars represent the standard deviation of replicates (n=3).

Temperature-dependent mass spectra reveal complex dissociation for Bn-B* in both solvents, with subtle differences in charge state (**Figure 2D & 2E**). Bn-B* in D_2_O consistently populates lower average charge states for the complex and individual proteins across the full temperature range (25-85 °C). These charge state differences indicate reduced solvent-accessible surface area (SASA) relative to H_2_O.^40, 41^ Complex abundance analysis shows a statistically significant increase in the melting temperature in D_2_O by 3 °C, consistent with enhanced thermal stability that aligns with solution melting results (**Figures 2F** and S9). As Bn-B* dissociates, the charge states for the individual proteins increase, especially for Bn, indicating that a loss of structure is necessary for the loss of the femtomolar binding affinity.^20^ Prior studies on the individual protein’s thermal stability shows Bn has a lower apparent T_m_ while B* has a very similar T_m_ to the complex, indicating the final loss of complex is associated with Bn unfolding following the initial unfolding of B*.^18^ The mass spectra suggests that thermal stability of Bn-B* is also altered in D_2_O and the different charge distributions on the proteins, may suggest unequal contribution from the individual proteins.

### Conformation Resolved Ion Mobility Profiles and Population Shifts

Ion mobility profiles for each CI-2 charge state (5+ - 10+) reveal many coexisting conformations increase systematically in CCS with charge state in both solvents (**Figure 3A**). Across all charge states, D_2_O favors the smaller CCS conformations as can be seen with the changes in relative abundance, which suggests a decrease in SASA of CI-2 in D_2_O. Additional compact conformations can be clearly seen for the 9+ and 10+ charge states in D_2_O, suggesting some sort of alteration in intramolecular and/or kinetic traps induced by protein-solvent interactions.^42^ Gaussian deconvolution resolves all native (N), intermediate (I), and extended (E) states across the entire charge state range, enabling quantitative population analysis (see supporting Figures S3, S10 and Table S2). A distinct transition conformation is observed across the 7+ and 8+ charge states at approximately 1331 Å² (dashed line, **Figure 3A**), which closely coincides with the Rayleigh charge limit of CI-2 (6.81 charges) and marks the transition between folded and unfolded conformational ensembles.^43^ The presence of many conformations on either side of this transition highlights the complexity of the unfolding process, even for a model two-state folding protein such as CI-2.^12, 44^ Due to such conformational complexity and the scope of this paper focusing on the effect of solution environment, the analysis and presentation of the data is focused on the most dominant conformations in each charge state.

**Figure 3:**
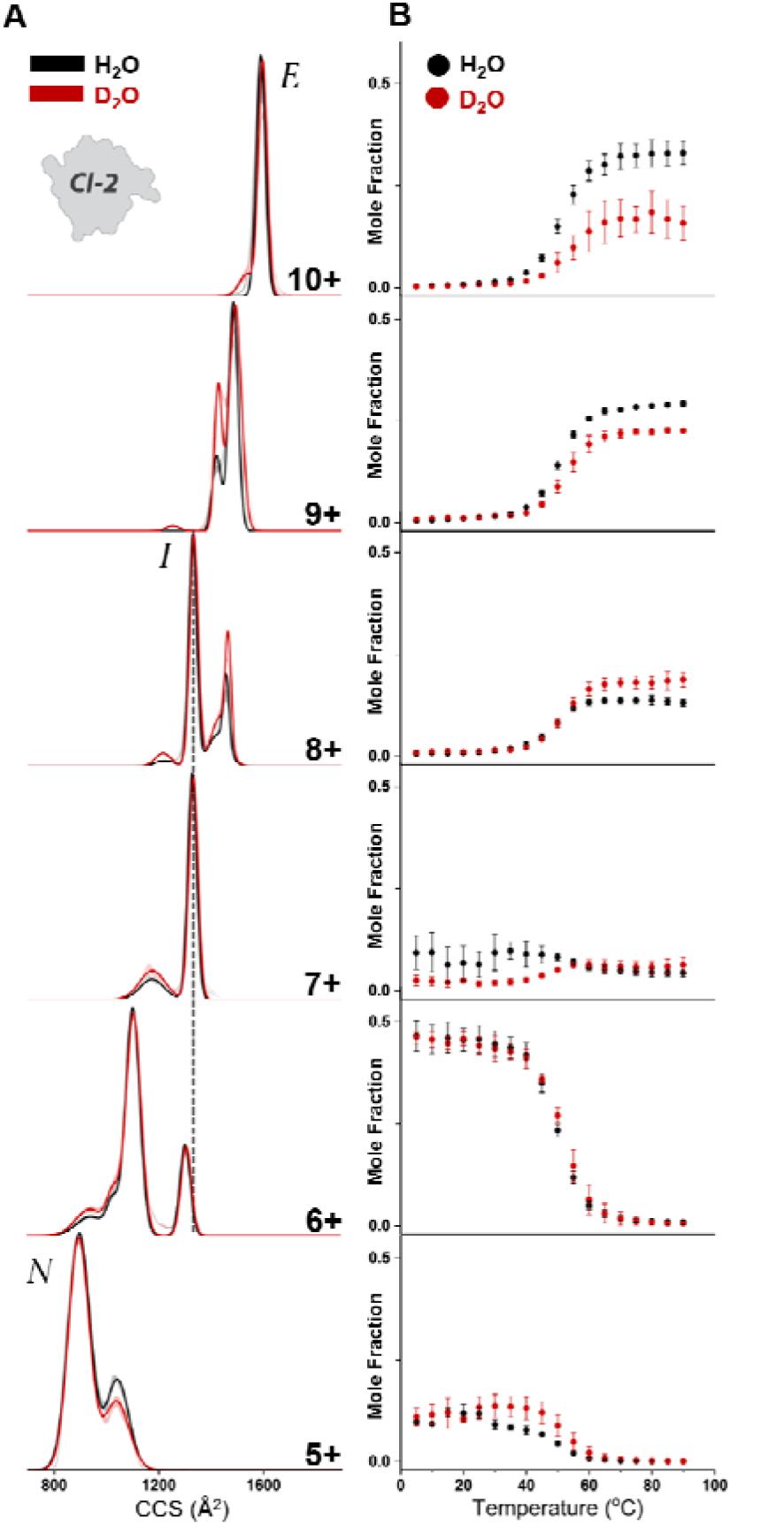
A) Averaged ion mobility profiles for each charge state 5+ - 10+, in H_2_O and D_2_O at temperatures when the charge state is most abundant for both H_2_O and D_2_O. As charge increases, the CCS of CI-2 increases and populates many conformations, where CI-2 tends to populate more compact conformations across all charge states in D_2_O. **B)** Mole fraction of the dominant conformation in each charge state across temperature in both H_2_O and D_2_O. The lower charge state (5+ & 6+) conformations are mostly similar, whereas the higher charge states (9+ & 10+) show significantly lower population of those conformations in D_2_O. Across all charge states, the inflection points (T_m_ or T_f_) are at higher temperatures in D_2_O (see **Table S3**), which aligns with the observed stability in D_2_O from the ensembl Z_avg_ data in Figure 2D. Fitting of conformations within each charge state to get the abundance is shown in supporting information Figure S3.

In every charge state of CI-2, the temperature at which native-like conformations decrease, and non-native conformations emerge is shifted to higher temperature in D_2_O relative to H₂O (**Figure 3B**). Sigmoidal fits to the abundance data yields higher inflection temperatures for D_2_O across all conformational families, demonstrating that solvent substitution globally stabilizes compact states and delays formation of extended conformations (Supporting Table S3). The dominant 7+ conformation is the outlier as it has opposing behavior (T_m_ in H_2_O but T_f_ in D_2_O), which dictates that the conformation requires more energy to become accessible in D_2_O. Notably, the stabilization observed in D_2_O is not limited to a single charge state but is distributed across the entire unfolding pathway, indicating a solvent-mediated effect on the conformational landscape rather than selective stabilization of a specific structural state.

Ion mobility profiles for each Bn-B* complex charge state (6+ - 10+) reveal a consistent increase in CCS and mobility broadening associated with an increase in SASA (**Figure 4A**). Unlike CI-2, Bn-B* has remarkably similar mobility profiles between solvents as complex. The higher charge states of the individual proteins (Bn & B*) have higher abundance for compact conformations in D_2_O as seen for CI-2 (see Figure S11), which we interpret are a result of exposed sites upon dissociation that hydration has a stronger influence on.^42^ Gaussian deconvolution for all conformers of the complex and individual proteins for quantitative population analysis can be found in supporting information (Figures S4). The 7+ to 10+ have increasingly broader CCS profiles with a similar lower CCS bound, indicating structural rearrangement that occurs to populate the higher charge states prior to dissociation of the complex.^43^ As for the temperature dependence of the dominant conformations in each charge state, the difference arises in the T_m_ and abundance (**Figure 4B**). D_2_O favors a higher abundance of the lower charge states (7-8+), which decreases in abundance to populate the 9-10+ charge states prior to complex dissociation. The highest charge states clearly show the T_m_ of the conformations are also delayed in D_2_O, indicating some interplay between conformation and complex thermal stability.

**Figure 4:**
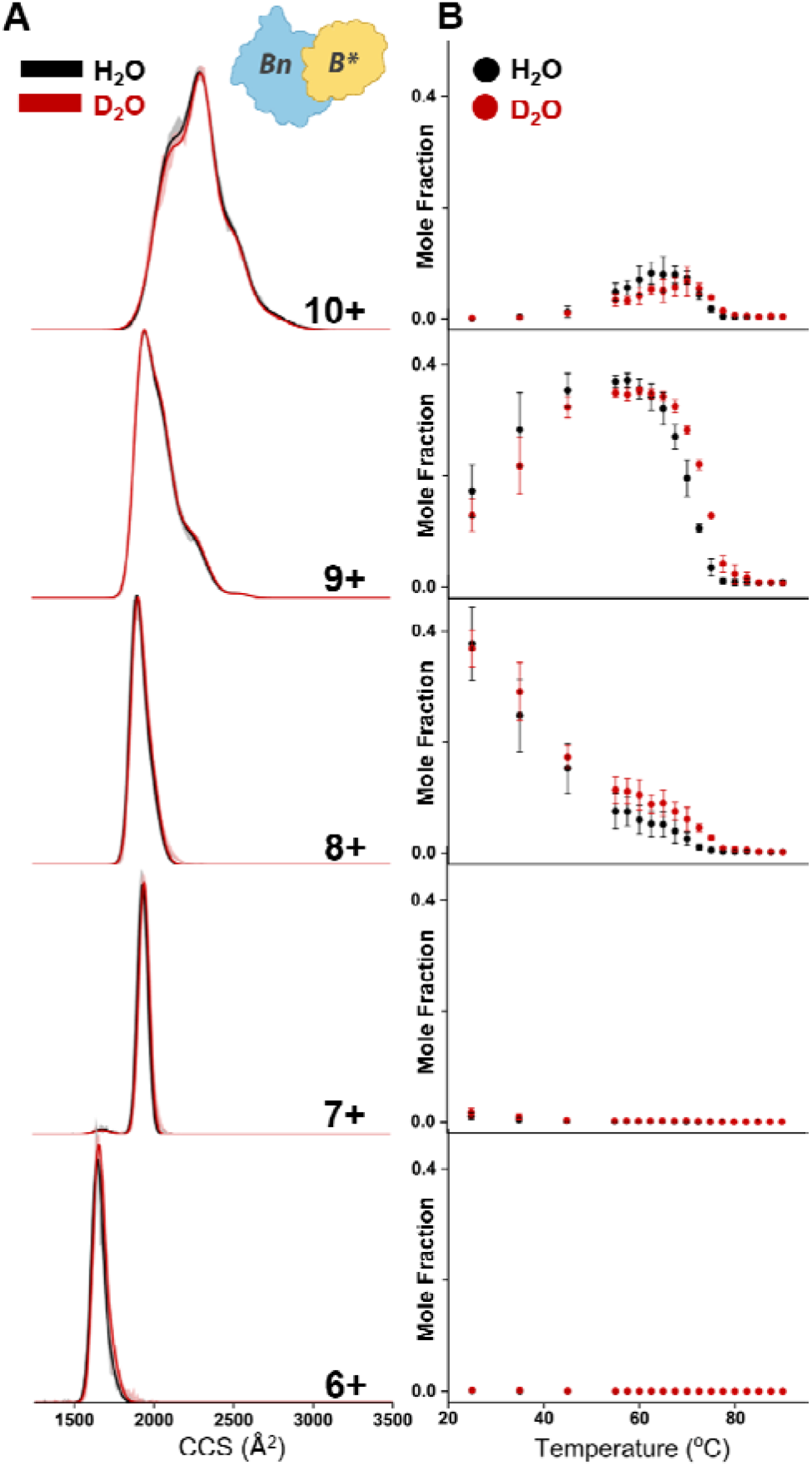
A) Averaged ion mobility profiles for each charge state 6+ - 10+, in H_2_O and D_2_O at temperatures when the charge state is most abundant for both H_2_O and D_2_O. As charge increases, the CCS of Bn-B* increases an distributions become broader. **B)** Mole fraction of the dominant conformation in each charge state across temperature in both H_2_O and D_2_O. The lower charge states (7+ - 8+) see overall slightly higher abundance in D_2_O, whereas th higher charge states (9+ - 10+) are slightly higher in H_2_O. Across all charge states, the inflection points (T_m_ or T_f_) ar at higher temperatures in D_2_O (see **Table S4**), which aligns with the observed stability in D_2_O from the ensemble complex stability data in Figure 2F. Fitting of conformations within each charge state to get the abundance is shown in supporting information Figure S4.

### Conformation Specific Thermodynamic Analysis

Using the conformation-resolved thermodynamic workflow (**Figure 1**), Gibbs free energies (ΔG) were extracted for all resolved conformations across temperature. At low temperature (20 °C), compact low-charge conformations of CI-2 exhibit favorable free energies (−ΔG), while extended high-charge conformations are unfavorable (+ΔG) in both solvents (**Figure 5A**). However, native-like conformations in D_2_O show systematically more favorable ΔG values than in H_2_O, indicating enhanced thermodynamic stability. At the melting regime (∼50 °C), nearly all conformations become thermodynamically accessible in both solvents (**Figure 5B**), reflecting coexistence of native and non-native conformations. Even in this regime, native states in D_2_O retain slightly more favorable ΔG values, consistent with delayed unfolding and higher T_m_. At high temperature (80 °C), extended conformations dominate in both solvents; however, these unfolded states are less thermodynamically favorable in D_2_O than in H_2_O (**Figures 5C** and S12). These data provide evidence that there are shallower free-energy wells for unfolded conformations in D_2_O relative to H_2_O, which is likely due to changes in protein intramolecular interactions perturbed by protein hydration ^45, 46^

**Figure 5:**
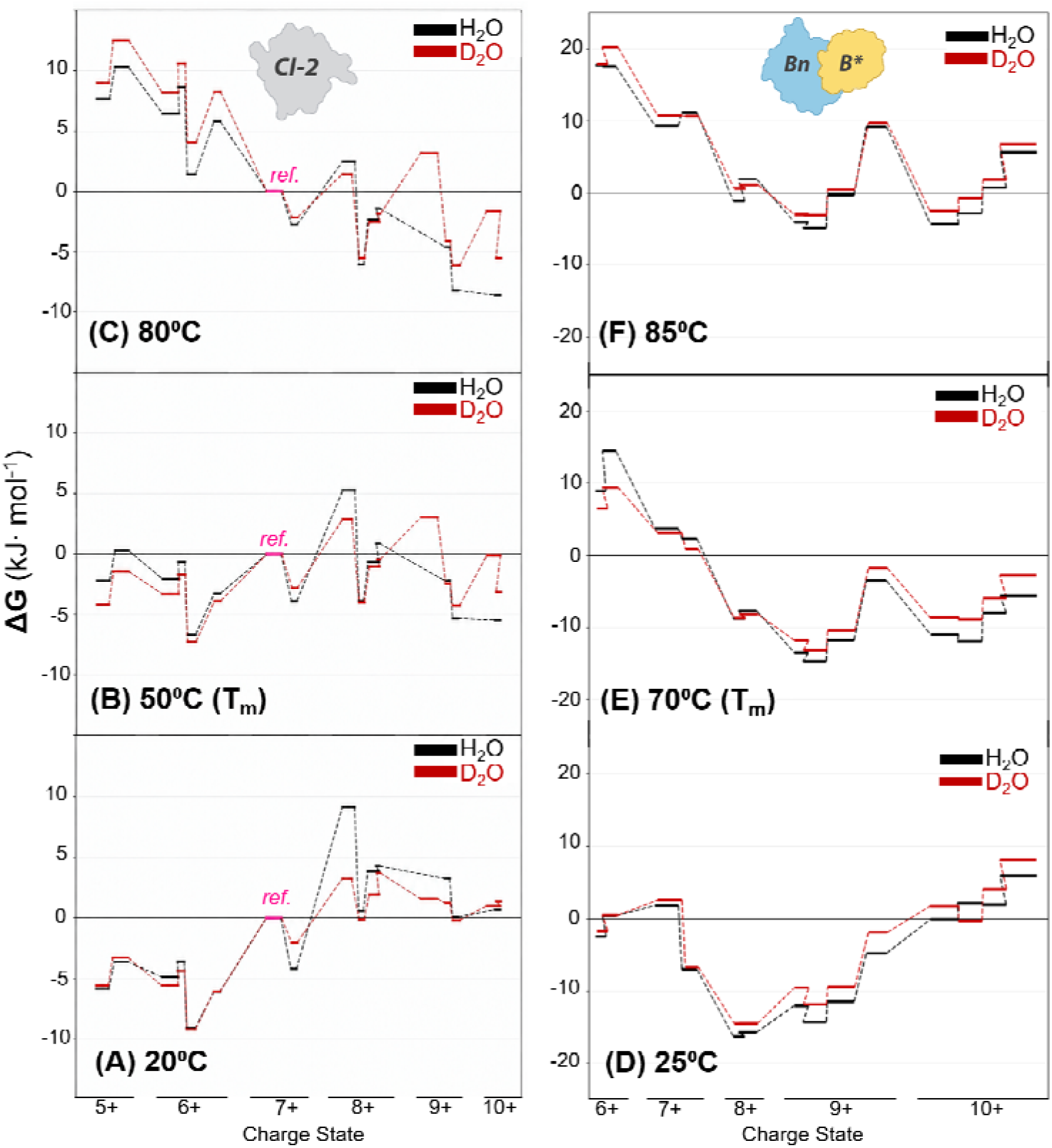
Favorability (ΔG) of each conformation in either solvent H_2_O (black) and D_2_O (red) separated by charge state and then CCS at **A)** 20□C, **B)** 50□C, and **C)** 80□C for CI-2. At 20□C, the lower charge states (smaller CCS) have favorable ΔG while the higher charge states (larger CCS) are not favorable. Around the melting temperature at 50 C almost all conformations are favorable in both solvents which aligns with native and non-native states being in comparable abundances. At high temperatures only the higher charge states are favorable, and it is important to not that in D_2_O the higher charge states do not have as favorable ΔG as H_2_O. Conformer favorability (ΔG) of Bn-B* conformations are seen in **D)** 25□C, **E)** 70□C, and **F)** 85□C for both solvents. At lower temperatures conformations are favorable for lower charge states, but as the complex gets to the melting temperature, the higher charge states dominate. At the highest temperature, most states are not favored or have low favorability due to their low abundanc in solution. Between solvents there are subtle differences in favorability for low charge states (6-8+), however at all temperatures, D_2_O has a more positive ΔG for the higher charge states (9-10+) indicating the protein does not as readily populate those larger SASA conformations.

A similar trend is seen for Bn-B* favorability where most conformations in the low charge states (6-8+) are more favorable (−ΔG) at low temperature (25 °C) in both solvents (**Figure 5D**). At the melting temperature (70 °C), nearly all conformations become thermodynamically accessible in both solvents (**Figure 5E**), with the lowest charge conformations becoming the most unfavored. At the highest temperature, most states are not favored (+ΔG) or have low favorability due to their low abundance in solution (**Figure 5F**). Between solvents there are subtle differences in favorability for low charge states (6-8+), however at all temperatures, D_2_O has a more positive ΔG for the higher charge states (9-10+) indicating the protein complex does not as readil populate those larger SASA conformations.

Van’t Hoff analysis (Figure S13) yields enthalpic (ΔH) and entropic (−TΔS) contributions for each conformation across all temperatures. At each of CI-2 conformation’s respective melting or formation temperatures (**Figure 6A**) compact, low-charge conformations (N) exhibit favorable enthalpy (−ΔH), consistent with stabilizing intramolecular interactions, whereas extended conformations (E) show unfavorable enthalpy (+ΔH) but favorable entropy (−TΔS), reflecting increased conformational dynamics and solvent-protein interactions.^47^ Analysis of the data by combining the conformational data into an “ensemble” set as seen in supporting Figure S14 shows that there is again a higher melting temperature of the CI-2 in D_2_O by ∼3 C (p-value = 0.04), more positive enthalpy and a less positive heat capacity (ΔC_p, H2O_ = 3.7 kJ·mol^-1^·K^-1^ and ΔC_p, D2O_ = 1.42 kJ·mol^-1^·K^-1^), suggesting there are more compact conformations present in D_2_O, which aligns with the IM-MS data. Notably, the ensemble results agree closely with prior work done by Jackson *et. al.* using scanning microcalorimetry, supporting this workflow a complementary high resolution approach to traditional solution-phase unfolding measurements.^12^ Additionally, the discrete clustering of H_2_O and D_2_O datasets demonstrate solvent-specific thermodynamic signatures for each conformation, despite similar compensation behavior.^48, 49^ Across the entire conformational landscape, the D_2_O conformations have more favorable enthalpy, which suggests there are more intramolecular contacts than in H_2_O. Notably the non-native states are less entropically favored in D_2_O, which provides evidence that they are less hydrated and/or occupy less microstates.^50, 51^ This separation indicates that while the fundamental unfolding mechanism is conserved, the solution environment alters the balance of enthalpic and entropic contributions, consistent with changes in protein-solvent interactions.

**Figure 6:**
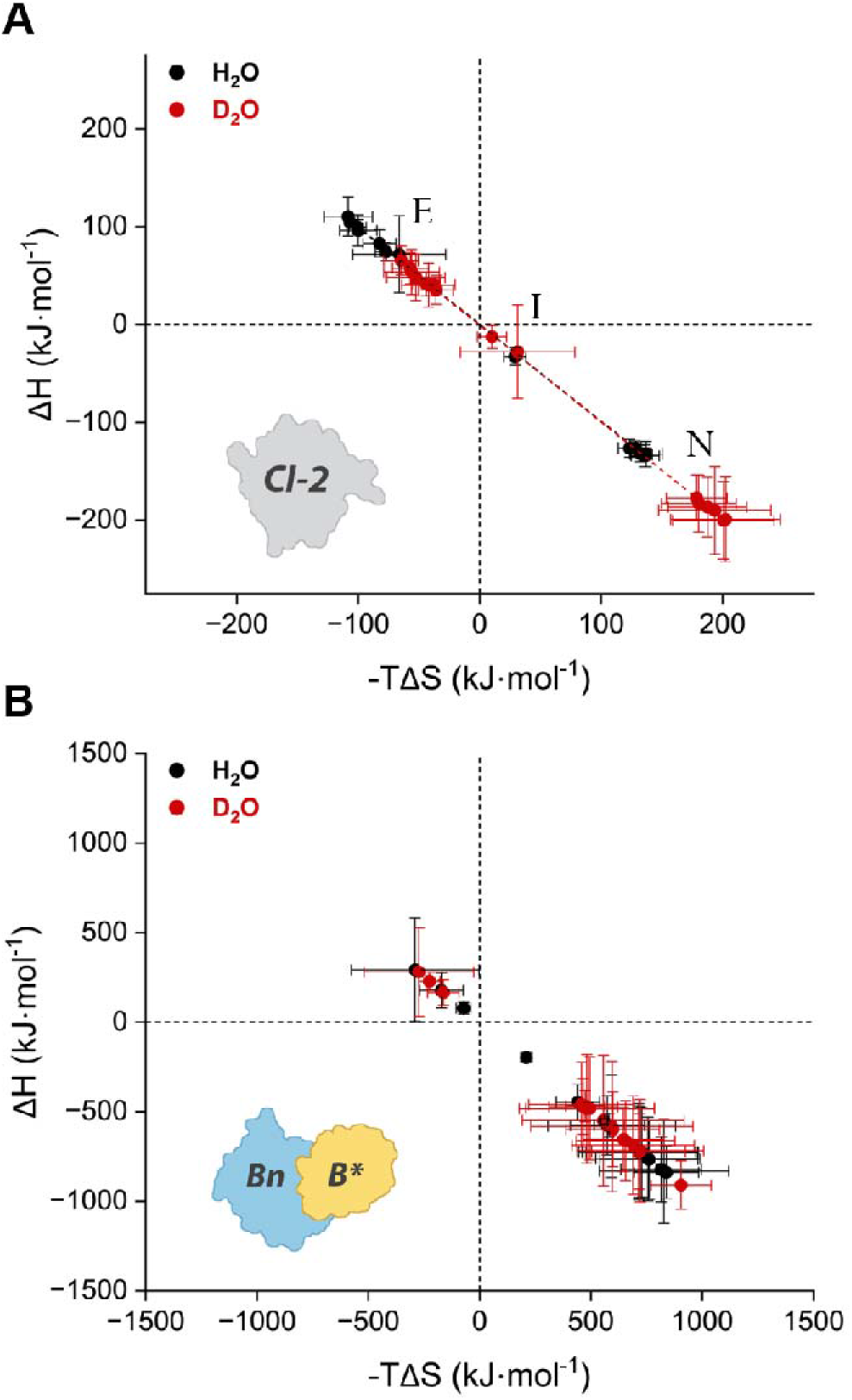
Enthalpy and entropy of each conformation at their respective melting or formation temperature for **A)** CI-2 and **B)** Bn-B* complex. The slopes in both solvents and protein systems are very close to -1, indicating strong entropy enthalpy compensation (EEC) which represents a similar response to the changes in hydration induced by temperature. However, for CI-2 the discreet localization of points separates the different types of conformations present along the unfolding pathway: native (N), intermediate (I) and extended (E). More favorable enthalpic contribution (-ΔH) of each conformation in D_2_O (red) indicates there are more intramolecular interactions present and hydration (protein-solvent interactions) of the conformations are altered. This CI-2 data additionally indicates that the more extended conformations have unfavorable enthalpy (+ΔH), but favorable entropy (-TΔS), indicating less structure. Interestingly, for Bn-B*, there is less favorable enthalpic contribution (-ΔH) of each conformation in D_2_O. When looking at the individual protein’s EEC plots (see **Figure S14**), D_2_O has an opposite effect on their thermodynamic signatures. For Bn, there is less entropy for the conformations in D_2_O, however for B* there is more entropy for each conformation. All together the decreased enthalpy of the complex is likely a result of the competition between thermodynamic signatures for the individual proteins (Figure S15). This data may also indicate the proteins have an unequal contribution to the complex thermal stability, with B* contributing the most. Error bars represent standard deviation of replicates (n=3).

Strong entropy enthalpy compensation (EEC) is also observed in both solvents, with near-unity slopes in ΔH versus −TΔS plots for Bn-B* as well (**Figure 6B**).^49^ Conformations from the 6-10+ charge state are favored enthalpically (+ΔH), while the very low abundant conformations for the 12+ charge state are favored entropically (−TΔS). Interestingly the opposite trend is seen where there is overall less favorable enthalpy for the complex in D_2_O. EEC plots of the individual proteins yield opposite trends for the effects of D_2_O (Figure S15), where Bn conformations are favored more enthalpically like in the case of CI-2, while B* is favored more entropically. This data may suggest that the complex conformational thermodynamics are unequally contributed from the individual proteins. Prior mutagenesis literature has been shown that B* residue D40 mutation to alanine causes the largest increase in K_d_, potentially supporting the larger role that B* has in the thermodynamics of the complex.^19^ Additionally, the relatively pronounced error bars seen in **Figure 6**, although smaller than the measured values of ΔH and -TΔS are related to the values being reported at the systems highest level of dynamics (T_m_).^52, 53^

### Differences in Protein-Solvent Interactions

Collision induced unfolding (CIU) experiments were performed on hydrogenated and deuterated CI-2 and Bn-B* to isolate solvent effects from intrinsic isotope labeling. CIU directly measures the enthalpic strength of the protein in the absence of solvent.^32^ Any change in CIU behavior between hydrogenated and perdeuterated CI-2 would indicate deuterium isotopes on the protein are affecting intramolecular bond strengths as suggested by Haidar *et. al*.^7^ IM-MS activation profiles and unfolding curves for the two charge states of both proteins are indistinguishable between isotopologues, and CIU_50_ values show no statistically significant difference (supporting Figures S16 and S17). This equivalence demonstrates that deuterium incorporation into the protein does not alter the unfolding behavior seen in **Figure 2**. When combined with the solvent-dependent thermal unfolding data, this directly implicates solvent-mediated interactions as the dominant source of stabilization, rather than isotope-dependent changes in protein-protein intramolecular hydrogen bond strength.^9^

To further investigate the influence of protein-solvent interactions, MDS of CI-2 were completed in pure H_2_O and D_2_O to analyze the structures and interactions globally and at the residue level. **Figure 7A** shows that CI-2 across the entire trajectory has a lower SASA in D_2_O by ∼3 nm^2^ on average, which is consistent with our observations in IM-MS where we see a preference for smaller CCS conformations (**Figures 3A and S18**). Looking at the SASA of each residue (Figure S19) the difference between solvents arises mainly around the dynamic loop region between the two beta sheets 40V-48R, where the charged residues appear to prefer more solvent exposure in D_2_O, but the nonpolar residues prefer to be less solvent exposed. Across the rest of the protein sequence, the beta sheet region and c-terminus show slightly smaller FWHM and SASA for most residues in D_2_O, which supports the overall protein SASA being smaller in D_2_O.

**Figure 7:**
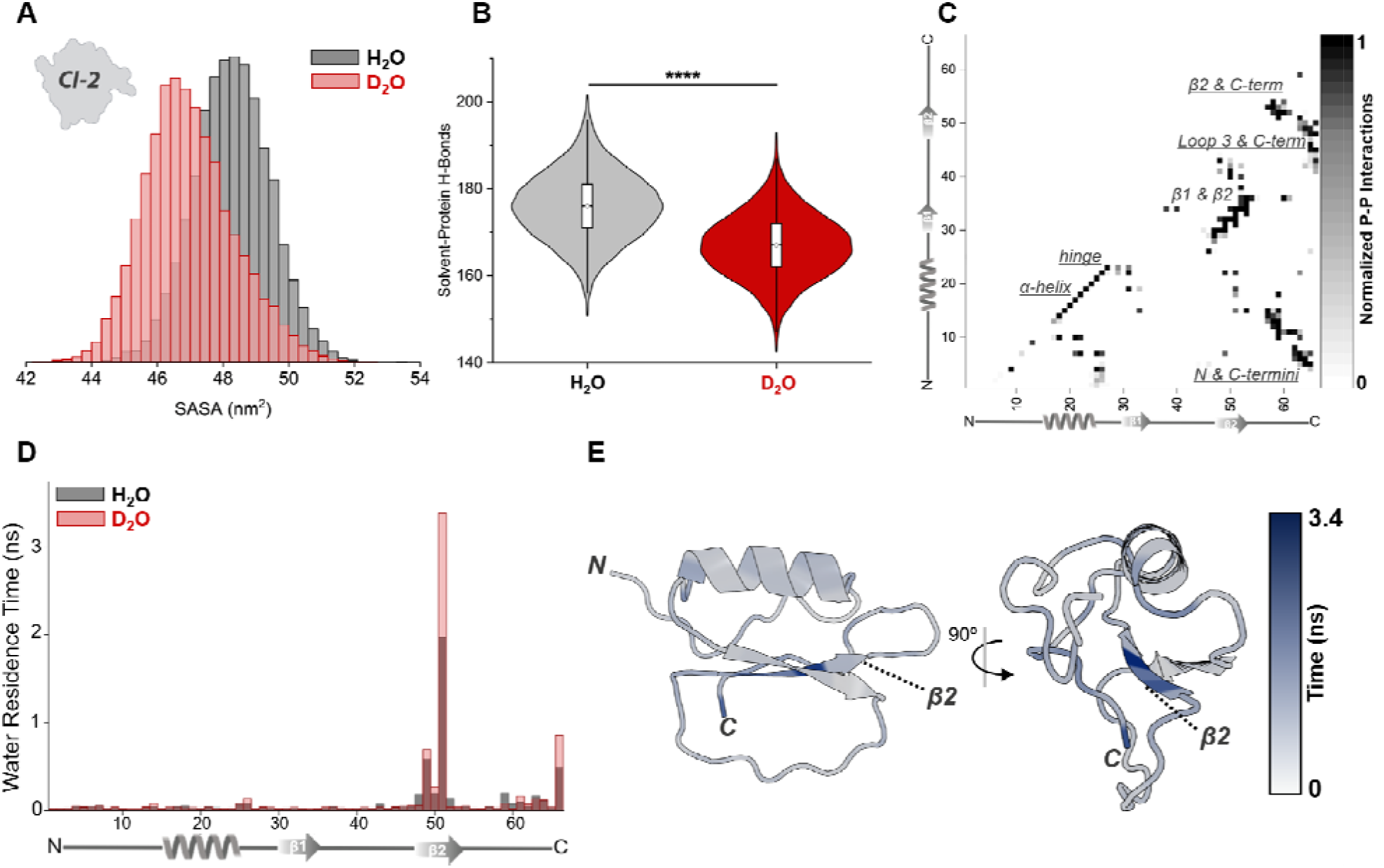
A) SASA of CI-2 in both H_2_O and D_2_O from each frame of the entire simulation. **B)** Violin plot showing the solvent-protein H-bonds for the entire protein across each frame of the molecular dynamic simulation in H_2_O an D_2_O. The average # of protein-solvent H-bonds are 175.9 ± 7.9 for H_2_O and 166.8 ± 7.4 for D_2_O with n=20000 and p-value <0.0001. These results indicate CI-2 in deuterated solvents have statistically less solvent-protein interactions which contributes to alterations in protein conformation and unfolding behavior. **C)** Protein-protein contact map that aligns with secondary structure and indicates the C-terminal end of the protein contains the most interactions and is thus most likely to contribute to the protein’s stability. **D)** Water residence time for each residue between solvent’s which shows the water interacts with the second β-sheet and C-terminus more often in D_2_O. **E)** Representative structure with the water residence time mapped onto it.

Sampling the number of solvent-protein hydrogen bonds across the entire MDS trajectory as seen in **Figure 7B**, there is a statistically significant decrease in solvent-protein hydrogen bonds for CI-2 in D_2_O (H_2_O ∼176 vs D_2_O ∼167). At the residue level (Figure S20), the results are similar to the SASA data in Figure S19, with the most different region being the dynamic loop region 40V-48R and the C-terminus. Looking at the various protein-protein contacts (**Figure 7C**), it is clear that the dominant number of stabilizing interactions are focused around the beta-sheet 2 and C-terminus. Electron-capture dissociation data (Figure S21) corroborates with the beta sheet region being the last region to unfold as CI-2 populates higher charge states, making it reasonable to conclude the dynamic loop region to the C-terminus is also highly important for the increased abundance for compact conformations in D_2_O. Water residence time analysis results in **Figure 7D**, aligns with the protein-contact map, where the regions with largest number of protein-contacts also has the longest residence times. Although there are fewer overall numbers of solvent-protein hydrogen bonds observed in D_2_O, water-residence time analysis shows that there are substantially longer residence times at the beta sheet 2 and C-terminus regions. **Figure 7E**, maps the water residence time onto a representative structure of CI-2, clearly indicating the localization of this effect to the beta sheet 2 and C-terminus region and defining the possibility that this region is most likely a dominant region that contributes to the different thermal stability observed in D_2_O.

MDS of Bn-B* was completed in a similar manner to CI-2 to compare the effects of solvent (H_2_O vs D_2_O) on the complex interactions. **Figure 8A** shows a less dynamic and lower SASA in D_2_O on average. The bimodal distribution in H_2_O correlates precisely with the bimodal SASA distribution seen for Bn (See Figure S22), which indicates that Bn is substantially stabilized in D_2_O. **Figure 8B** again shows a statistical less amount of solvent-protein hydrogen bonds in D_2_O with the largest difference in SASA between solvents being the reduced FWHM of residue SASA in D_2_O. The Bn-B* complex is well known to be facilitated by charged-residue interactions (salt bridges) and the protein contact map in **Figure 8C** articulates the negatively charged region of B* that interacts at the mostly positive enzymatic domain and regions on the same plane of Bn.^20, 21^ Again, water-residence times are the longest at the region with the most contacts (interface) **Figures 8D & 8E**, which implicates water interactions being important in the interface as shown in prior work.^54, 55^ Additionally, D_2_O promotes longer water-residence times than H_2_O at the complex interface (**Figures 8D**), which suggests the waters may be involved in the altered thermal stability of the complex.

**Figure 8:**
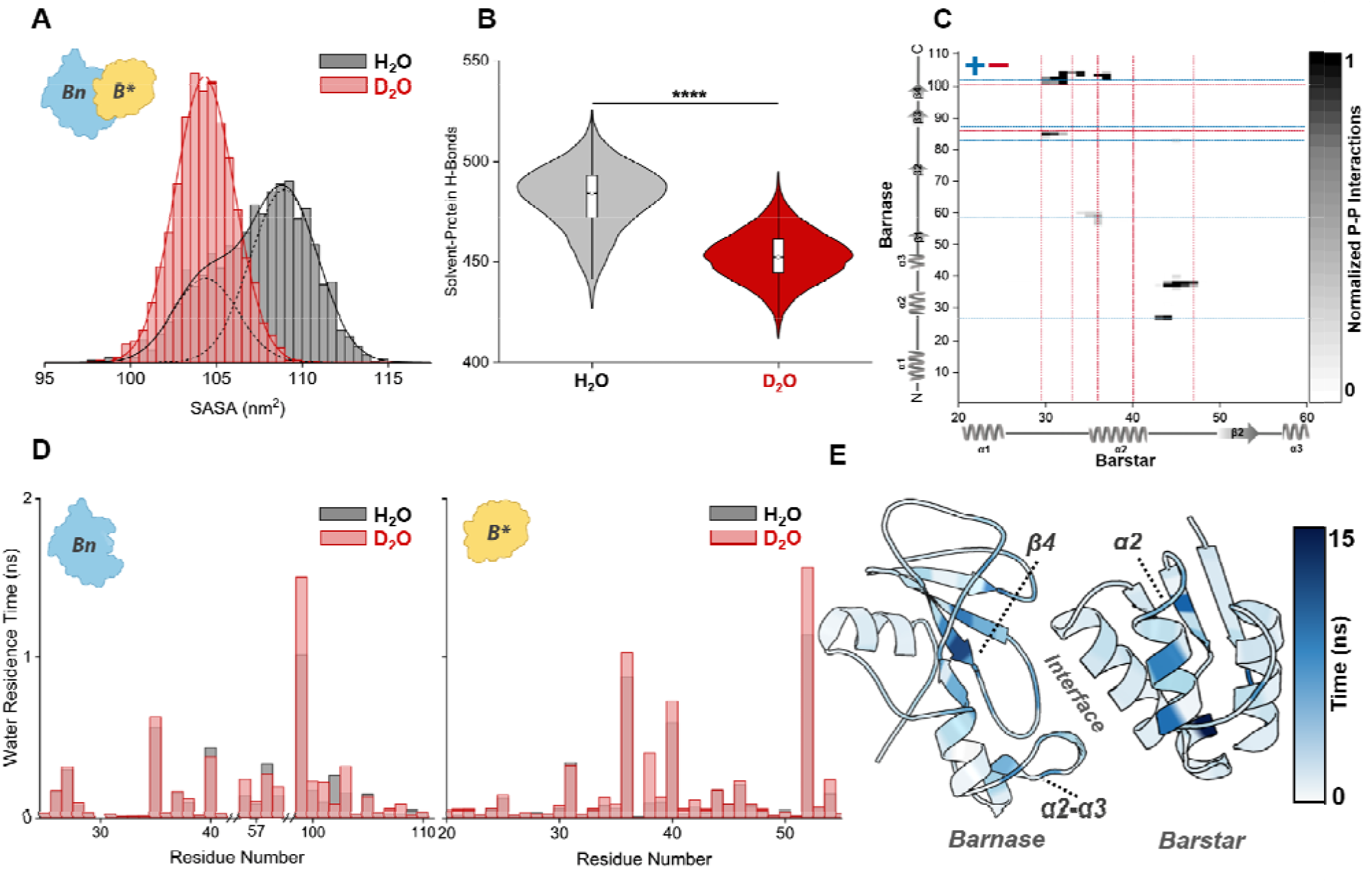
A) SASA of Bn-B* in both H_2_O and D_2_O from each frame of the entire simulation. The bimodal distributio of the complex in H_2_O is a result of contributions from Bn. **B)** Violin plot showing the solvent-protein H-bonds for the entire protein across each frame of the molecular dynamic simulation in H_2_O and D_2_O. These results indicate Bn-B* in deuterated solvents have statistically less solvent-protein interactions. **C)** Protein-protein contact map that aligns with secondary structure and indicates regions of both proteins that contribute to the complex. Red and blue lines indicate negative and positive residues that form the complex interface. **D)** Water residence time for each residue that makes up the complex interface. Across almost all residues D_2_O again has longer residence times that may contribute to the increased complex stability. **E)** Representative structures with the interface turned 45° towards the reader with the water residence time mapped onto it show that the region with the longest water-residence time is the interface, implicating water-interactions as an important part of the Bn-B* stability.

## Discussion

A common stability signature emerges from two different protein systems. CI-2 and Bn-B* differ in size, topology, and dominant non-covalent interactions.^12, 21^ CI-2 undergoes unfolding, whereas Bn-B* loses the assembled complex as the temperature increases. Nonetheless, D_2_O produces four common signatures: an approximately 3 °C increase in the ensemble melting temperature, reduced favorability of extended conformations, lower global protein hydration and location specific longer water residence times. The agreement across systems allows for the discussion of whether solvent and/or isotope mediated interactions contribute to thermal stability and more broadly to the altered dynamics proteins are observed to have in D_2_O.^8, 37, 39, 56^

The first common signature is the approximately 3 °C increase in ensemble melting temperature in D_2_O (**Figure 2**). This magnitude of stabilization is consistent with the solvent effect reported for other proteins, where D_2_O commonly increases the thermal transition temperature relative to H_2_O. ^7^ The usual interpretation is that D_2_O is a less favorable solvent for the exposed, nonpolar surface generated during unfolding, increasing the hydrophobic contribution to compact-state stability due to stronger water-water hydrogen bonding.^57^ In this work, D_2_O does not necessarily create fundamentally different structures; instead, it changes the energetic balance between the compact and extended populations (**Figure 5**).^58^

The second common observation is the reduced favorability of extended conformations in D_2_O. In CI-2, this is observed as less favorable higher charge conformations at elevated temperatures (**Figure 3)**. For Bn-B*, **Figures 2D,E** and **4** show higher charge-state conformations that precede complex dissociation are also consistently less populated in D_2_O. The calculated free energy of these extended conformations across temperatures in the two solvents again supports their decreased favorability (**Figure 5**). These observations suggest that the main effect of D_2_O is not simply to strengthen the native basin, but to reduce the accessibility of conformations with increased SASA.^5, 59^ The free-energy landscape for CI-2 in supporting information Figure S23 visualizes this behavior, where the unfolded highest charge states have shallower wells in D_2_O and are therefore less favorable to populate.^51^ Additionally, the thermodynamic signatures of conformations have stronger enthalpy for CI-2 in D_2_O, which is consistent with prior literature while there is less enthalpy in D_2_O for the complex (**Figure 6**).^60, 61^ B* conformations were also observed to have less enthalpy in D_2_O, which along with prior mutagenesis work and structures (Figures S15 & S24) positions water mediated interactions to be one of the dominant interactions that govern the complex stability.^19^ These results again support the notion that D_2_O does not always act on proteins in the same manner.^37^ The favorability interpretation also agrees with the lower average charge-states and more compact mobility distributions in D_2_O, indicating that the solvent affects the whole conformational ensemble rather than only the native or extended conformations.

The third is reduced global protein hydration in D_2_O. Both simulations show lower average SASA and fewer protein-solvent hydrogen bonds in D_2_O (**Figure 7 & 8**), while the IM-MS results independently show a preference for lower charge-state and more compact conformations (**Figure 2-4**). MD simulations and experiments have reported that proteins can become more compact and rigid in D_2_O and that fewer solvent molecules participate in hydrogen bonds with the protein relative to H_2_O. Reduced hydration can preserve intramolecular contacts by limiting solvent competition with hydrogen-bonding and other local interactions.^5, 62^ In H_2_O, more frequent protein-water interactions can facilitate local disruption of these contacts and lower the energetic cost of moving from compact states into intermediate, unfolded or dissociation-prone conformations.^46^ In contrast, the reduced extent of hydration in D_2_O may limit solvent-driven perturbation of native contacts, increasing the energetic barriers for escape from the folded basin (Figure S23).^2^ Thus, reduced global hydration provides a useful explanation for why D_2_O favors compact states, but it does not by itself identify which protein regions control the stabilization.

The last common feature is the increase in location-specific water residence time in D_2_O. Although fewer total protein-solvent hydrogen bonds are present in D_2_O, the waters that remain at key sites persist longer. For CI-2, the longest residence times are localized at β-sheet 2 and the C-terminal region, which contain the highest density of intramolecular contacts and remain structured late in the unfolding pathway (**Figure 7** and S21). For Bn-B*, the longest residence times are localized at the protein-protein interface, where direct charged contacts and water-mediated interactions maintain the complex (**Figure 8**).^15^ This observation reconciles the apparently opposing trends of lower global hydration and greater stability: D_2_O does not simply dehydrate the protein surface, but instead also reorganizes hydration around structurally important regions. ^5^ A smaller number of longer-lived waters may stabilize local interactions by reducing fluctuations, or facilitating residue interactions.^17, 45, 46, 63^ It is important to note, this study cannot define whether the longer residence times are due to simple hydration kinetics (viscosity), however it seems more likely that the lower zero-point vibrational energy of deuterium is increasing the water hydrogen bond strengths and thus residence times.^9^ It is to our understanding that the consideration of site-specific hydration is not typically considered as a contributor to different biomolecular behavior in D_2_O, although plenty of examples exist within the literature of important-water motifs for function and binding.^5, 64^

Together, these signatures support the conclusion that D_2_O stabilization arises from changes in global and site specific hydration interactions. The CIU data are important in this regard because hydrogenated and deuterated proteins show indistinguishable intramolecular unfolding behavior, indicating that deuterium incorporation into the protein is not sufficient to explain the thermal shifts. Instead, D_2_O appears to reduce global protein hydration while increasing the lifetime of water interactions at sites that are particularly important for structural stability. For CI-2, these sites are concentrated in the second β-sheet/C-terminal region; for Bn-B*, they are concentrated at the binding interface. This site-specific residence-time effect provides a molecular basis for the altered conformational dynamics in D_2_O and suggests that water residence time may be a useful metric for identifying hydration sites that disproportionately contribute to protein folding and protein-protein complex stability.^46, 65^ More broadly, these results demonstrate that solvent identity can alter protein free-energy landscapes by tuning solvent-protein interactions, which in turn can affect the stability and dynamics of proteins.^37–39, 66^

## Conclusion

The effects of D_2_O on protein physicochemical behavior are system dependent motivating a closer examination of how solvent-dependent hydration influences certain biophysical properties and interactions. Here, we show that the increased thermal stability of proteins in D_2_O is associated with altered hydration dynamics, not with deuterium exchange into the protein.^8^ CI-2 and Bn-B* represent very different thermal processes: unfolding of a small single-domain protein and loss of a picomolar affinity protein complex. Yet both show the same ∼ 3 °C increased melting temperatures, increased compact-state populations, and reduced accessibility of the extended conformations required for unfolding or dissociation in D_2_O. The CIU data further demonstrate that these differences are not retained after removal of solvent, supporting a solvent-mediated origin for the thermal stability change observed.^7^

The central result is that D_2_O changes hydration in two ways. Globally, both proteins have lower SASA and fewer solvent-protein hydrogen bonds, consistent with reduced solvent competition for intramolecular and interfacial interactions. However, water remains longer at sites/regions that matter most for stability. In CI-2, these sites are concentrated at the β-sheet 2/C-terminal structural core and in Bn-B*, they occur at the charged binding interface. Therefore, D_2_O does not simply produce less hydrated proteins. It also produces longer-lived hydration at the structural regions that provide protein stability.^63^ Future work with MDS or by other means may elucidate the thermodynamic contribution of these longer-lived/stronger hydrogen bonded waters on the stability of proteins.^17^ Combining site-specific mutagenesis and thermodynamic thermal stability assays in light and heavy water may provide additional evidence to support our claims and/or another contributing factor to the increased thermal stability of proteins in D_2_O.^12, 19^ More broadly, we believe that there is a need to incorporate hydration in most discussions to fully understand a protein’s physicochemical properties and the methods used herein are some of the few capable of gleaming insight on the phenomenon.^67, 68^

## Supporting Information

Supporting Information includes detailed instrument parameters, thorough examination of the thermodynamic workflow, conformational fitting, and MDS results.

## Acknowledgement

Funding for this project was provided by the National Institutes of Health (Grants R01GM138863 and RM1GM149374), the Robert A. Welch Foundation (grant A-2162-20230405), and endowment funds from an MDS SCIEX Professorship. Computing resources were provided by the Texas A&M High-Performance Research Computing center. We gratefully acknowledge Professor A. Josh Wand for providing the barnase-barstar plasmid used in this work.

## Author Contributions

The manuscript was written through contributions of all authors. All authors have given approval to the final version of the manuscript.

## References

(1) Galano-Frutos, J. J.; Sancho, J. Energy, water, and protein folding: A molecular dynamics-based quantitative inventory of molecular interactions and forces that make proteins stable. Protein Science 2024, 33 (2). DOI: 10.1002/pro.4905.

(2) Schiffer, C. A.; Dötsch, V. The role of protein-solvent interactions in protein unfolding. Current Opinion in Biotechnology 1996, 7 (4), 428–432. DOI: 10.1016/s0958-1669(96)80119-4.

(3) El-Baba, T. J.; Fuller, D. R.; Hales, D. A.; Russell, D. H.; Clemmer, D. E. Solvent Mediation of Peptide Conformations: Polyproline Structures in Water, Methanol, Ethanol, and 1-Propanol as Determined by Ion Mobility Spectrometry-Mass Spectrometry. Journal of the American Society for Mass Spectrometry 2019, 30 (1), 77–84. DOI: 10.1007/s13361-018-2034-7.

(4) Pan, H.; Raab, S. A.; El-Baba, T. J.; Schrecke, S. R.; Laganowsky, A.; Russell, D. H.; Clemmer, D. E. Variation of CI-2 Conformers upon Addition of Methanol to Water: An IMS-MS-Based Thermodynamic Analysis. The Journal of Physical Chemistry A 2023, 127 (45), 9399–9408. DOI: 10.1021/acs.jpca.3c03651.

(5) Spyrakis, F.; Ahmed, M. H.; Bayden, A. S.; Cozzini, P.; Mozzarelli, A.; Kellogg, G. E. The Roles of Water in the Protein Matrix: A Largely Untapped Resource for Drug Discovery. Journal of Medicinal Chemistry 2017, 60 (16), 6781–6827. DOI: 10.1021/acs.jmedchem.7b00057.

(6) Bellissent-Funel, M.-C.; Hassanali, A.; Havenith, M.; Henchman, R.; Pohl, P.; Sterpone, F.; van der Spoel, D.; Xu, Y.; Garcia, A. E. Water Determines the Structure and Dynamics of Proteins. Chemical Reviews 2016, 116 (13), 7673–7697. DOI: 10.1021/acs.chemrev.5b00664.

(7) Haidar, Y.; Konermann, L. Effects of Hydrogen/Deuterium Exchange on Protein Stability in Solution and in the Gas Phase. Journal of the American Society for Mass Spectrometry 2023, 34 (7), 1447–1458. DOI: 10.1021/jasms.3c00130.

(8) Giubertoni, G.; Bonn, M.; Woutersen, S. D2O as an Imperfect Replacement for H2O: Problem or Opportunity for Protein Research? The Journal of Physical Chemistry B 2023, 127 (38), 8086–8094. DOI: 10.1021/acs.jpcb.3c04385.

(9) Clark, T.; Heske, J.; Kühne, T. D. Opposing Electronic and Nuclear Quantum Effects on Hydrogen Bonds in H2O and D2O. ChemPhysChem 2019, 20 (19), 2461–2465. DOI: 10.1002/cphc.201900839.

(10) Martini, S.; Bonechi, C.; Foletti, A.; Rossi, C. Water-Protein Interactions: The Secret of Protein Dynamics. The Scientific World Journal 2013, 2013 (1), 1–6. DOI: 10.1155/2013/138916.

(11) Crilly, C. J.; Eicher, J. E.; Warmuth, O.; Atkin, J. M.; Pielak, G. J. Water’s Variable Role in Protein Stability Uncovered by Liquid-Observed Vapor Exchange NMR. Biochemistry 2021, 60 (41), 3041–3045. DOI: 10.1021/acs.biochem.1c00552.

(12) Jackson, S. E.; Fersht, A. R. Folding of chymotrypsin inhibitor 2. 1. Evidence for a two-state transition. Biochemistry 1991, 30 (43), 10428–10435. DOI: 10.1021/bi00107a010.

(13) Gazi, R.; Maity, S.; Jana, M. Thermal stability and hydration dynamics of Chymotrypsin Inhibitor 2 in aqueous amino acid solutions from replica exchange molecular dynamics. Biochemical and Biophysical Research Communications 2026, 806, 153403. DOI: 10.1016/j.bbrc.2026.153403.

(14) Raab, S. A.; El-Baba, T. J.; Woodall, D. W.; Liu, W.; Liu, Y.; Baird, Z.; Hales, D. A.; Laganowsky, A.; Russell, D. H.; Clemmer, D. E. Evidence for Many Unique Solution Structures for Chymotrypsin Inhibitor 2: A Thermodynamic Perspective Derived from vT-ESI-IMS-MS Measurements. Journal of the American Chemical Society 2020, 142 (41), 17372–17383. DOI: 10.1021/jacs.0c05365.

(15) Schreiber, G.; Fersht, A. R. Interaction of barnase with its polypeptide inhibitor barstar studied by protein engineering. Biochemistry 1993, 32 (19), 5145–5150. DOI: 10.1021/bi00070a025.

(16) Frisch, C.; Schreiber, G.; Johnson, C. M.; Fersht, A. R. Thermodynamics of the interaction of barnase and barstar: changes in free energy versus changes in enthalpy on mutation 1 1Edited by J. Karn. Journal of Molecular Biology 1997, 267 (3), 696–706. DOI: 10.1006/jmbi.1997.0892.

(17) Chong, S.-H.; Ham, S. Dynamics of Hydration Water Plays a Key Role in Determining the Binding Thermodynamics of Protein Complexes. Scientific Reports 2017, 7 (1). DOI: 10.1038/s41598-017-09466-w.

(18) Makarov, A. A.; Protasevich, I. I.; Lobachov, V. M.; Kirpichnikov, M. P.; Yakovlev, G. I.; Gilli, R. M.; Briand, C. M.; Hartley, R. W. Thermostability of the barnase—barstar complex. FEBS Letters 1994, 354 (3), 251–254. DOI: 10.1016/0014-5793(94)01127-3.

(19) Schreiber, G.; Fersht, A. R. Energetics of protein-protein interactions: Analysis ofthe Barnase-Barstar interface by single mutations and double mutant cycles. Journal of Molecular Biology 1995, 248 (2), 478–486. DOI: 10.1016/s0022-2836(95)80064-6.

(20) Tidor, B.; Lee, L.-P. Barstar is electrostatically optimized for tight binding to barnase. Nature Structural Biology 2001, 8 (1), 73–76. DOI: 10.1038/83082.

(21) Lee, L. P.; Tidor, B. Optimization of binding electrostatics: Charge complementarity in the barnase-barstar protein complex. Protein Science 2001, 10 (2), 362–377. DOI: 10.1110/ps.40001.

(22) Caro, J. A.; Valentine, K. G.; Cole, T. R.; Wand, A. J. Pressure, motion, and conformational entropy in molecular recognition by proteins. Biophysical Reports 2023, 3 (1), 100098. DOI: 10.1016/j.bpr.2022.100098.

(23) Holmberg, A.; Blomstergren, A.; Nord, O.; Lukacs, M.; Lundeberg, J.; Uhlén, M. The biotin-streptavidin interaction can be reversibly broken using water at elevated temperatures. ELECTROPHORESIS 2005, 26 (3), 501–510. DOI: 10.1002/elps.200410070.

(24) Ayan, E.; Yuksel, B.; Destan, E.; Ertem, F. B.; Yildirim, G.; Eren, M.; Yefanov, O. M.; Barty, A.; Tolstikova, A.; Ketawala, G. K.;, et al. Cooperative allostery and structural dynamics of streptavidin at cryogenic- and ambient-temperature. Communications Biology 2022, 5 (1). DOI: 10.1038/s42003-021-02903-7.

(25) Kr żel, A.; Bal, W. A formula for correlating p K a values determined in D 2 O and H 2 O. Journal of Inorganic Biochemistry 2004, 98 (1), 161–166. DOI: 10.1016/j.jinorgbio.2003.10.001.

(26) McCabe, J. W.; Mallis, C. S.; Kocurek, K. I.; Poltash, M. L.; Shirzadeh, M.; Hebert, M. J.; Fan, L.; Walker, T. E.; Zheng, X.; Jiang, T.;, et al. First-Principles Collision Cross Section Measurements of Large Proteins and Protein Complexes. Anal Chem 2020, 92 (16), 11155–11163. DOI: 10.1021/acs.analchem.0c01285 From NLM Medline.

(27) Chen, S.-H.; Russell, D. H. How Closely Related Are Conformations of Protein Ions Sampled by IM-MS to Native Solution Structures? Journal of the American Society for Mass Spectrometry 2015, 26 (9), 1433–1443. DOI: 10.1007/s13361-015-1191-1.

(28) Ruotolo, B. T.; Benesch, J. L. P.; Sandercock, A. M.; Hyung, S.-J.; Robinson, C. V. Ion mobility–mass spectrometry analysis of large protein complexes. Nature Protocols 2008, 3 (7), 1139–1152. DOI: 10.1038/nprot.2008.78.

(29) Van’T Hoff, M. J. H. Etudes de dynamique chimique. Recueil des Travaux Chimiques des Pays-Bas 1884, 3 (10), 333–336. DOI: 10.1002/recl.18840031003.

(30) Seelig, J.; Seelig, A. Protein Unfolding—Thermodynamic Perspectives and Unfolding Models. International Journal of Molecular Sciences 2023, 24 (6), 5457. DOI: 10.3390/ijms24065457.

(31) Streicher, W. W.; Makhatadze, G. I. Unfolding Thermodynamics of Trp-Cage, a 20 Residue Miniprotein, Studied by Differential Scanning Calorimetry and Circular Dichroism Spectroscopy. Biochemistry 2007, 46 (10), 2876–2880. DOI: 10.1021/bi602424x.

(32) Jeon, C. K.; Rojas Ramirez, C.; Makey, D. M.; Kurulugama, R. T.; Ruotolo, B. T. CIUSuite 3: Next-Generation CCS Calibration and Automated Data Analysis Tools for Gas-Phase Protein Unfolding Data. Journal of the American Society for Mass Spectrometry 2024, 35 (8), 1865–1874. DOI: 10.1021/jasms.4c00176.

(33) Abraham, M. J.; Murtola, T.; Schulz, R.; Páll, S.; Smith, J. C.; Hess, B.; Lindahl, E. GROMACS: High performance molecular simulations through multi-level parallelism from laptops to supercomputers. SoftwareX 2015, 1–2, 19–25. DOI: 10.1016/j.softx.2015.06.001.

(34) Huang, J.; Mackerell, A. D. CHARMM36 all-atom additive protein force field: Validation based on comparison to NMR data. Journal of Computational Chemistry 2013, 34 (25), 2135–2145. DOI: 10.1002/jcc.23354.

(35) Xi, Z.; Kuo, S.-T.; Cong, X.; Yan, X.; Russell, D. H. Revealing Anisotropic Growth of Liraglutide Oligomers by Native Ion Mobility Mass Spectrometry and Molecular Dynamics Simulation. ACS Central Science 2025, 11 (7), 1154–1165. DOI: 10.1021/acscentsci.5c00431.

(36) Marklund, E. G.; Robinson, C. V.; Baldwin, A. J.; Benesch, J. L. P. Collision Cross Sections for Structural Proteomics. Structure 2015, 23 (4), 791–799. DOI: 10.1016/j.str.2015.02.010.

(37) Lantz, C.; Rider, R. L.; Kuo, S.-T.; Xi, Z.; Burningham, E.; Yun, S. D.; Laganowsky, A.; Russell, D. H. Hydration (H2O and D2O) Dictates the Stabilities and Conformational Entropy of Transthyretin Tetramers. Journal of the American Chemical Society 2025, 147 (39), 35616–35626. DOI: 10.1021/jacs.5c11055.

(38) Hampton, J.; Lantz, C.; Rider, R. L.; Yun, S. D.; Laganowsky, A.; Russell, D. H. Effects of Hydration on Transthyretin Conformational Dynamics and Oligomerization. Biochemistry 2025, 64 (23), 4571–4580. DOI: 10.1021/acs.biochem.5c00589.

(39) Evans, K. A.; Sun, H. M.; Powers, M.; Lantz, C.; Laganowsky, A.; Rye, H.; Russell, D. H. Roles of Hydration in Protein–Ligand Binding: Passive or Active Participant? The Journal of Physical Chemistry A 2025, 129 (40), 9477–9485. DOI: 10.1021/acs.jpca.5c04986.

(40) McCabe, J. W.; Shirzadeh, M.; Walker, T. E.; Lin, C. W.; Jones, B. J.; Wysocki, V. H.; Barondeau, D. P.; Clemmer, D. E.; Laganowsky, A.; Russell, D. H. Variable-Temperature Electrospray Ionization for Temperature-Dependent Folding/Refolding Reactions of Proteins and Ligand Binding. Anal Chem 2021, 93 (18), 6924–6931. DOI: 10.1021/acs.analchem.1c00870 From NLM Medline.

(41) Rider, R. L.; Lantz, C.; Fan, L.; Russell, D. H. Structure and Stabilities of Solution and Gas Phase Protein Complexes. Journal of the American Society for Mass Spectrometry 2024, 35 (12), 3028–3036. DOI: 10.1021/jasms.4c00306.

(42) Fort, K. L.; Silveira, J. A.; Pierson, N. A.; Servage, K. A.; Clemmer, D. E.; Russell, D. H. From Solution to the Gas Phase: Factors That Influence Kinetic Trapping of Substance P in the Gas Phase. The Journal of Physical Chemistry B 2014, 118 (49), 14336–14344. DOI: 10.1021/jp5103687.

(43) Susa, A. C.; Xia, Z.; Tang, H. Y. H.; Tainer, J. A.; Williams, E. R. Charging of Proteins in Native Mass Spectrometry. Journal of the American Society for Mass Spectrometry 2017, 28 (2), 332–340. DOI: 10.1007/s13361-016-1517-7.

(44) Christofi, E.; Barran, P. Ion Mobility Mass Spectrometry (IM-MS) for Structural Biology: Insights Gained by Measuring Mass, Charge, and Collision Cross Section. Chemical Reviews 2023, 123 (6), 2902–2949. DOI: 10.1021/acs.chemrev.2c00600.

(45) Cioni, P.; Strambini, G. B. Effect of heavy water on protein flexibility. Biophys J 2002, 82 (6), 3246–3253. DOI: 10.1016/S0006-3495(02)75666-X From NLM Medline.

(46) Guzzi, R.; Arcangeli, C.; Bizzarri, A. R. A molecular dynamics simulation study of the solvent isotope effect on copper plastocyanin. Biophysical Chemistry 1999, 82 (1), 9–22. DOI: 10.1016/s0301-4622(99)00097-6.

(47) Wand, A. J.; Sharp, K. A. Measuring Entropy in Molecular Recognition by Proteins. Annual Review of Biophysics 2018, 47 (1), 41–61. DOI: 10.1146/annurev-biophys-060414-034042.

(48) Dragan, A. I.; Read, C. M.; Crane-Robinson, C. Enthalpy-entropy compensation: the role of solvation. Eur Biophys J 2017, 46 (4), 301–308. DOI: 10.1007/s00249-016-1182-6 From NLM.

(49) Fox, J. M.; Zhao, M.; Fink, M. J.; Kang, K.; Whitesides, G. M. The Molecular Origin of Enthalpy/Entropy Compensation in Biomolecular Recognition. Annual Review of Biophysics 2018, 47 (1), 223–250. DOI: 10.1146/annurev-biophys-070816-033743.

(50) Kannan, A.; Naganathan, A. N. Ensemble origins and distance-dependence of long-range mutational effects in proteins. iScience 2022, 25 (10), 105181. DOI: 10.1016/j.isci.2022.105181.

(51) Dill, K. A.; Chan, H. S. From Levinthal to pathways to funnels. Nature Structural Biology 1997, 4 (1), 10–19. DOI: 10.1038/nsb0197-10.

(52) Sharp, K. Entropy—enthalpy compensation: Fact or artifact? Protein Science 2001, 10 (3), 661–667. DOI: 10.1110/ps.37801.

(53) Cooper, A. Protein Heat Capacity: An Anomaly that Maybe Never Was. The Journal of Physical Chemistry Letters 2010, 1 (22), 3298–3304. DOI: 10.1021/jz1012142.

(54) Chong, S. H.; Ham, S. Impact of chemical heterogeneity on protein self-assembly in water. Proc Natl Acad Sci U S A 2012, 109 (20), 7636–7641. DOI: 10.1073/pnas.1120646109 From NLM Medline.

(55) Copie, G.; Cleri, F.; Blossey, R.; Lensink, M. F. On the ability of molecular dynamics simulation and continuum electrostatics to treat interfacial water molecules in protein-protein complexes. Scientific Reports 2016, 6 (1), 38259. DOI: 10.1038/srep38259.

(56) Giubertoni, G.; Feng, L.; Klein, K.; Giannetti, G.; Rutten, L.; Choi, Y.; Van Der Net, A.; Castro-Linares, G.; Caporaletti, F.; Micha, D.;, et al. Elucidating the role of water in collagen self-assembly by isotopically modulating collagen hydration. Proceedings of the National Academy of Sciences 2024, 121 (11). DOI: 10.1073/pnas.2313162121.

(57) Sheu, S.-Y.; Schlag, E. W.; Selzle, H. L.; Yang, D.-Y. Molecular Dynamics of Hydrogen Bonds in Protein−D_2_O: The Solvent Isotope Effect. The Journal of Physical Chemistry A 2008, 112 (5), 797–802. DOI: 10.1021/jp0771668.

(58) Darby, J. F.; Hopkins, A. P.; Shimizu, S.; Roberts, S. M.; Brannigan, J. A.; Turkenburg, J. P.; Thomas, G. H.; Hubbard, R. E.; Fischer, M. Water Networks Can Determine the Affinity of Ligand Binding to Proteins. Journal of the American Chemical Society 2019, 141 (40), 15818–15826. DOI: 10.1021/jacs.9b06275.

(59) Ladbury, J. E. Just add water! The effect of water on the specificity ligand binding sites And its potential application in drug design. Chemistry and Biology 1996, 3, 973–980.

(60) Kaltashov, I. A.; Eyles, S. J. Crossing the phase boundary to study protein dynamics and function: combination of amide hydrogen exchange in solution and ion fragmentation in the gas phase. Journal of Mass Spectrometry 2002, 37 (6), 557–565. DOI: 10.1002/jms.338.

(61) Efimova, Y. M.; Haemers, S.; Wierczinski, B.; Norde, W.; Well, A. A. V. Stability of globular proteins in H2O and D2O. Biopolymers 2007, 85 (3), 264–273. DOI: 10.1002/bip.20645.

(62) Honegger, P.; Schmollngruber, M.; Steinhauser, O. Macromolecular crowding and the importance of proper hydration for the structure and dynamics of protein solutions. Physical Chemistry Chemical Physics 2018, 20 (29), 19581–19594. DOI: 10.1039/c8cp02360c.

(63) Caro, J. A.; Harpole, K. W.; Kasinath, V.; Lim, J.; Granja, J.; Valentine, K. G.; Sharp, K. A.; Wand, A. J. Entropy in molecular recognition by proteins. Proceedings of the National Academy of Sciences 2017, 114 (25), 6563–6568. DOI: 10.1073/pnas.1621154114.

(64) Russo, D.; Hura, G.; Head-Gordon, T. Hydration Dynamics Near a Model Protein Surface. Biophysical Journal 2004, 86 (3), 1852–1862. DOI: 10.1016/s0006-3495(04)74252-6.

(65) Luo, P.; Zhai, Y.; Senses, E.; Mamontov, E.; Xu, G.; Z, Y.; Faraone, A. Influence of Kosmotrope and Chaotrope Salts on Water Structural Relaxation. The Journal of Physical Chemistry Letters 2020, 11 (21), 8970–8975. DOI: 10.1021/acs.jpclett.0c02619.

(66) Sun, H. M.; Evans, K. A.; Powers, M.; Xi, Z.; Lantz, C.; Laganowsky, A.; Rye, H.; Russell, D. H. Allostery without Conformational Change: A Native Mass Spectrometry Perspective. The Journal of Physical Chemistry B 2025, 129 (34), 8668–8679. DOI: 10.1021/acs.jpcb.5c03261.

(67) Doan, L. C.; Dahanayake, J. N.; Mitchell-Koch, K. R.; Singh, A. K.; Vinh, N. Q. Probing Adaptation of Hydration and Protein Dynamics to Temperature. ACS Omega 2022, 7 (25), 22020–22031. DOI: 10.1021/acsomega.2c02843.

(68) Lee, A. L.; Sapienza, P. J. The Power of Protein Dynamics in Binding and Allostery. Biochemistry 2026, 65 (6), 649–660. DOI: 10.1021/acs.biochem.5c00782. TOC:

