## Supporting Information for "Site Specific Hydration Contributes to the Thermal Stability of Proteins and Protein-Complexes"

### Table of Contents


| | Page |
|---|---|
| Figure S1. | S2 |
| Figure S2. | S3 |
| Figure S3. | S4 |
| Figure S4. | S5 |
| Figure S5. | S6 |
| Figure S6. | S7 |
| Figure S7. | S8 |
| Figure S8. | S9 |
| Figure S9. | S10 |
| Figure S10. | S11 |
| Figure S11. | S12 |
| Figure S12. | S13 |
| Figure S13. | S14 |
| Figure S14. | S15 |
| Figure S15. | S16 |
| Figure S16. | S17 |
| Figure S17. | S18 |
| Figure S18. | S19 |
| Figure S19. | S20 |
| Figure S20. | S21 |
| Figure S21. | S22 |
| Figure S22. | S23 |
| Figure S23. | S24 |
| Figure S24. | S25 |
| Table S1. | S26 |
| Table S2. | S27 |
| Table S3. | S28 |
| Table S4. | S28 |
| Table S4. | S30 |
| References | S33 |

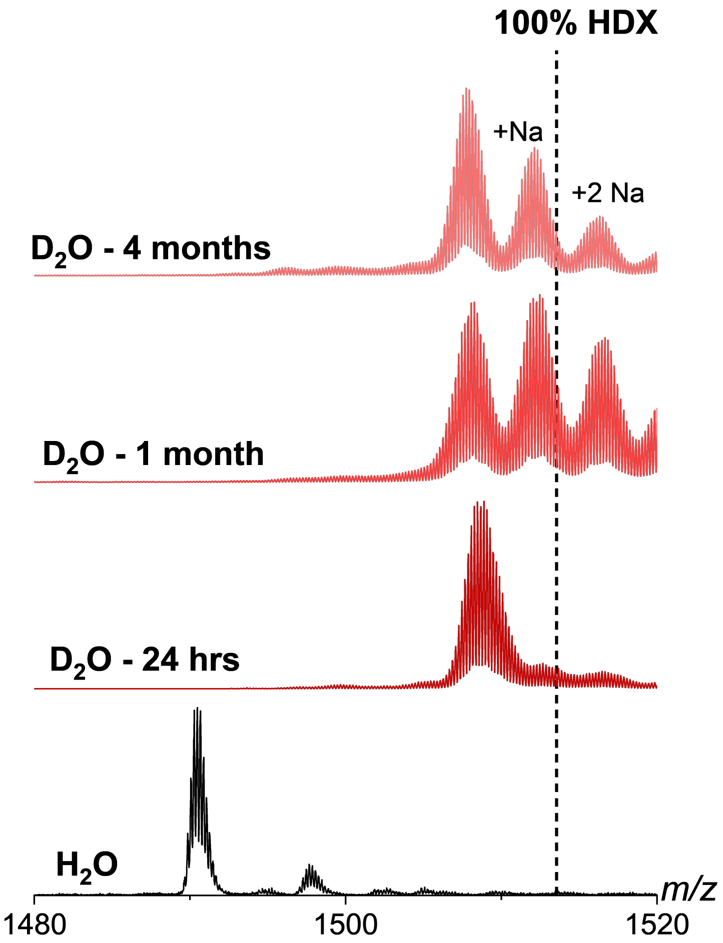


**Figure S1.** Mass differences for CI-2 in $H_2O$ and $D_2O$ with 10mM AA at 37 ºC across time observed with the 5+ charge state. There is no significant mass increase or reduction in peak FWHM from 24hrs to over 4 months incubation indicating denaturing CI-2 would be necessary to achieve closer 100% deuteration of titratable residues. Denaturing was not undertaken as it may perturb the native contacts and folding that are necessary to retain for comparison of protein unfolding between solvents. The maximum average HDX uptake was 79.0% at 24 h with a decline to 76.3% and 75.1% in 1 month and 4 months. The decline in deuterium over the long time periods we ascribe to changes in protein stability and/or interactions as evidence by the increase in sodium uptake.

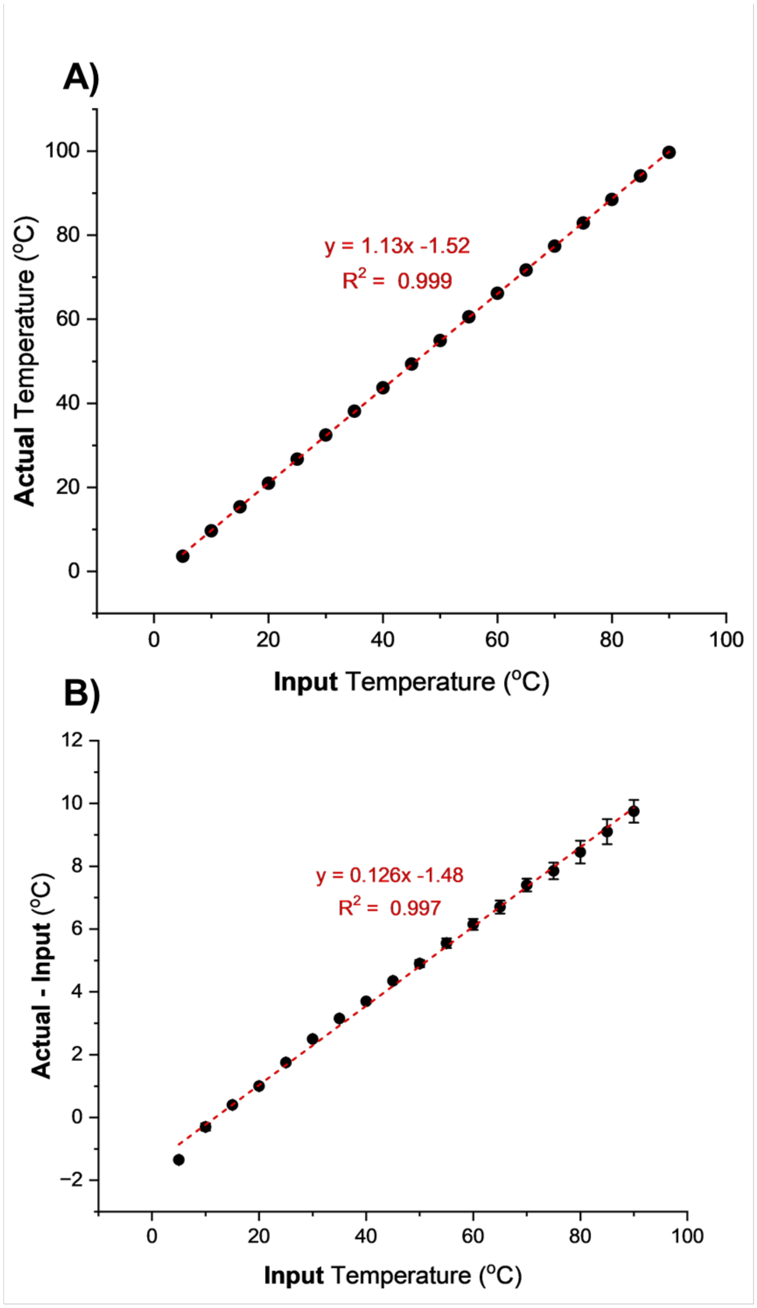


**Figure S2. A)** vT-ESI device calibration using an internal probe to measure the temperature of the solution in the nESI emitter (actual) versus the temperature input on the vT-ESI device software (input). **B)** Deviation of the actual solution temperature from the input showing a strong linear relation. The linear fit to the bottom plot (**B**) was used to calculate the needed input temperature to ensure the solution temperature was at the desired temperature (i.e. input = 63.9 ºC ; actual = 70.0 ºC). Error bars are present on both plots for all temperatures and represent standard deviation of replicates (n=3).

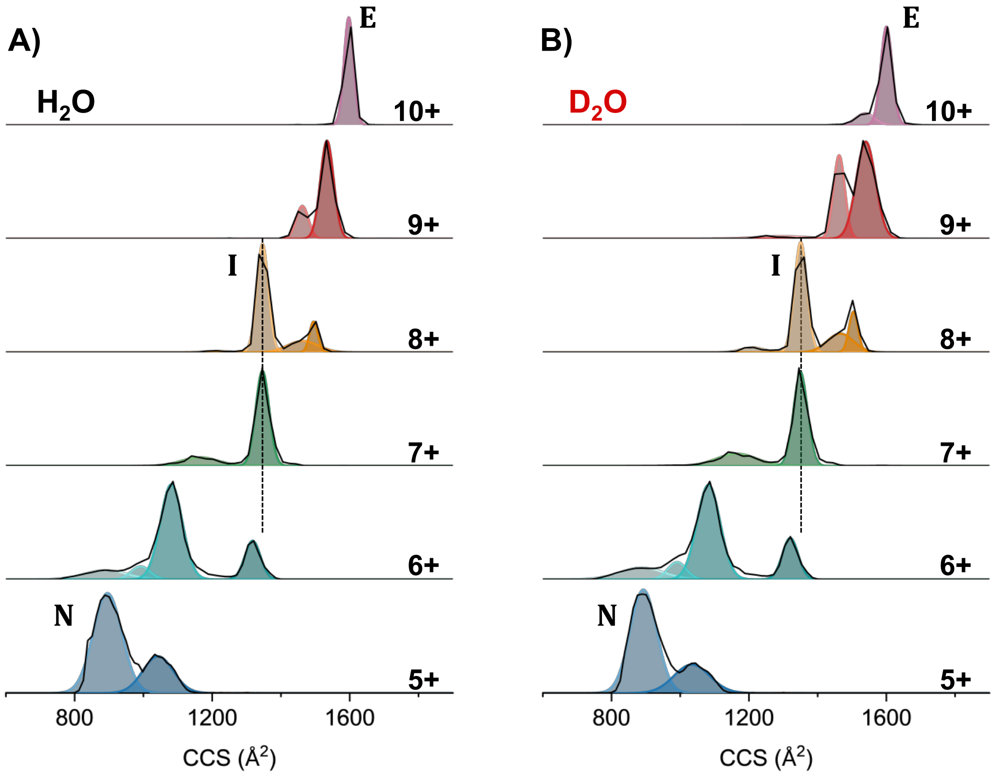


**Figure S3.** Representative raw IM-MS profiles for each CI-2 charge state (black lines) and their respective fits to resolve conformations in **A)** $H_2O$ and **B)** $D_2O$. CI-2 in $D_2O$ has a higher abundance of compact conformations across all charge states with the 9 & 10+ having the presence of smaller CCS conformations not seen in $H_2O$. Noted on both plots is the idea of native (N) intermediate (I) and extended (E) conformations that represent the unfolded transition measured however it is important to note the plethora of conformations between the N→E transition highlights how impactful IM-MS can be to gain high resolution information on protein unfolding.

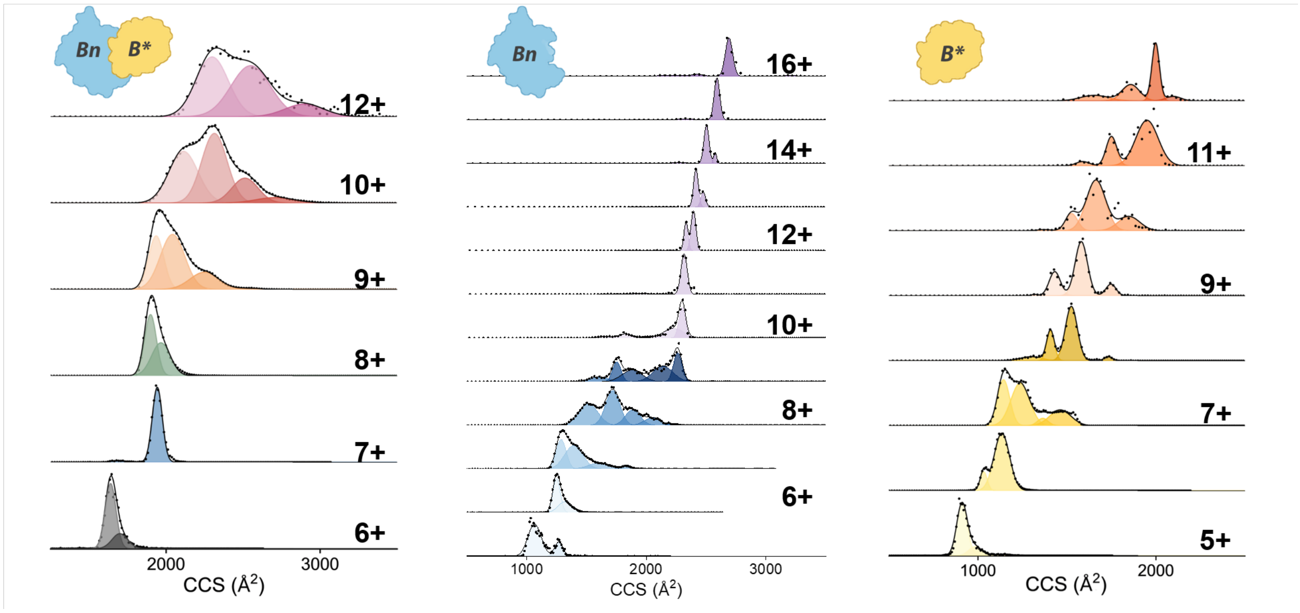


**Figure S4.** Representative raw IM-MS profiles for each Bn-B* charge state (black dots) and their respective fits to resolve conformations in $H_2O$. Notice the peak broadening for the complex and the narrowing of conformations for the individual proteins at higher charge states.

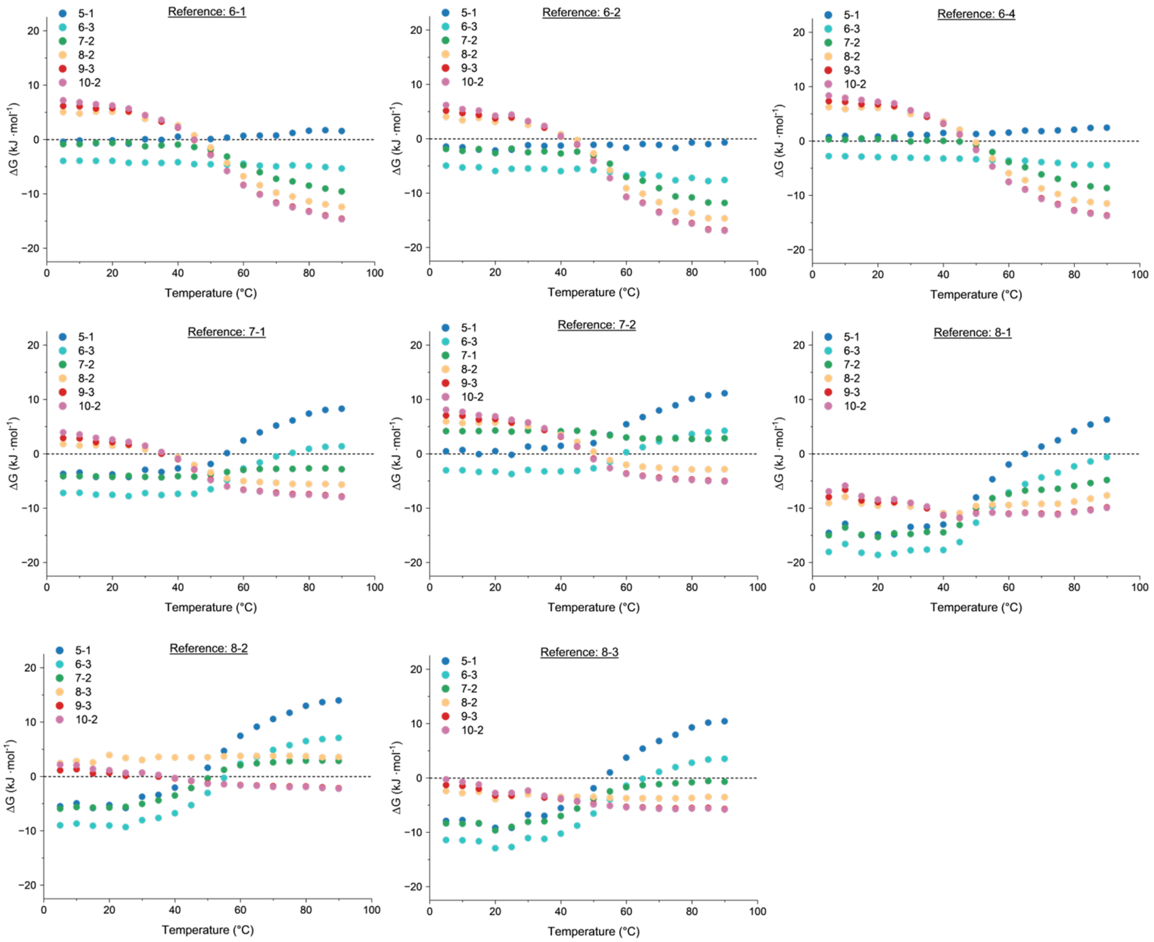


**Figure S5.** Comparison of the favorability (ΔG) of each dominant CI-2 conformation with different reference conformations chosen where the reference 7-1 conformation results are the most reflective of the charge state abundance changes. For this relative thermodynamic analysis it is imperative to choose a low abundance conformation within an intermediate charge state that remains abundant throughout the temperature range. Choosing otherwise gives results where certain conformations do not have clear changes in favorability (- to +) which is expected for the data to match the change of charge state and conformation abundance.

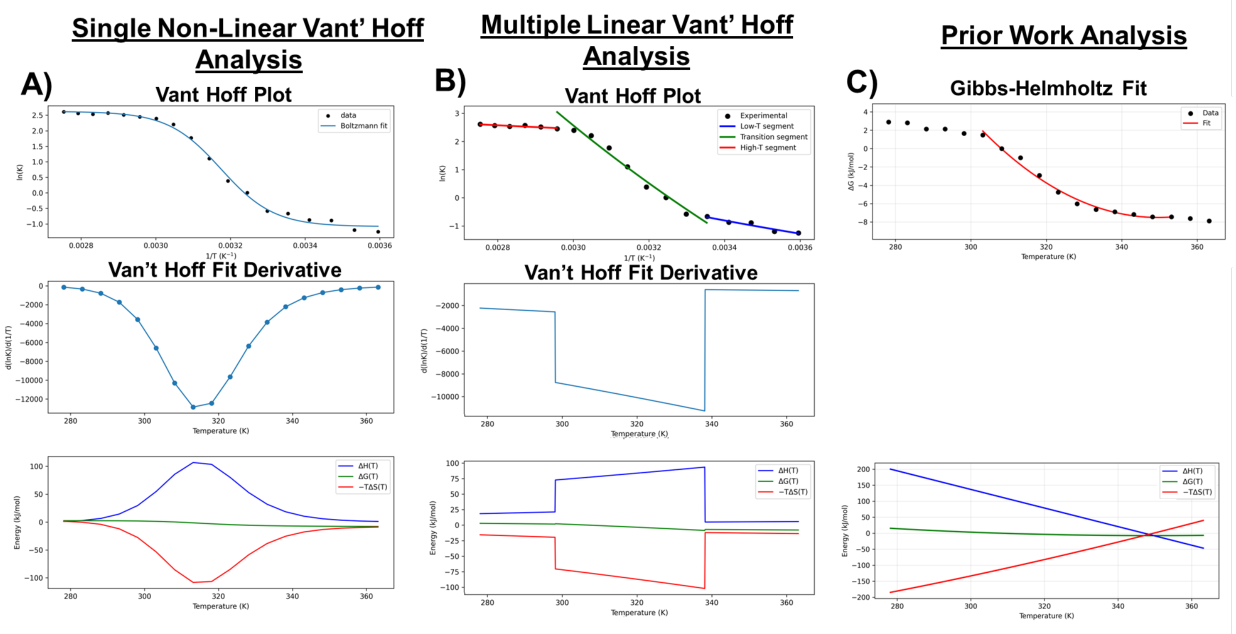


**Figure S6:** Comparison of different analysis for the most abundant conformation in the CI-2 9+ charge state (conformer 9-3). This work uses a single non-linear Van't Hoff workflow as seen in **A)** which can capture the experimental features/curvature of the data. This workflow as validated as the crude features of the thermodynamic data (ΔH and -TΔS) are captured by separating the data into multiple sections and analyzed using linear Van't Hoff analysis as seen in **B)**. **C)** Prior work uses the Gibbs Helmholtz equation to capture the inflection point really well but the limitation lies in it's ability to extract the data from the entire temperature when the data is no longer highly linear. At the midpoint of the data the following ΔH values were calculated to be **A)** 122.9 **B)** 81.3 **C)** 98.6 for each method respectively which shows their similarity at the midpoint to be on the same order of magnitude.

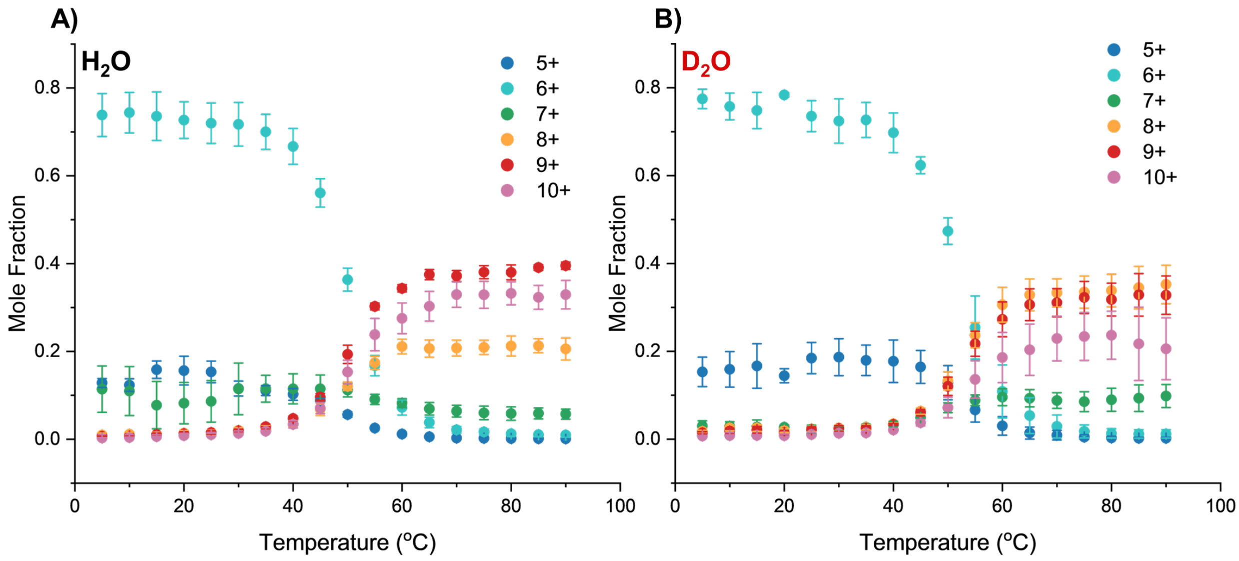


**Figure S7.** Comparison of the abundance of charge states as a function of temperature in **A)** $H_2O$ and **B)** $D_2O$. At low temperatures the 6+ dominates in both solutions with the abundance of the 5+ being more dominant in $D_2O$. At higher temperatures the 9 +and 10+ are most dominant in $H_2O$ while the 8+ and 9+ are dominant in $D_2O$. The higher abundance of lower charge states at both temperature extremes suggests that the SASA for CI-2 in $D_2O$ is lower. Error bars represent standard deviation of replicates (n=3).

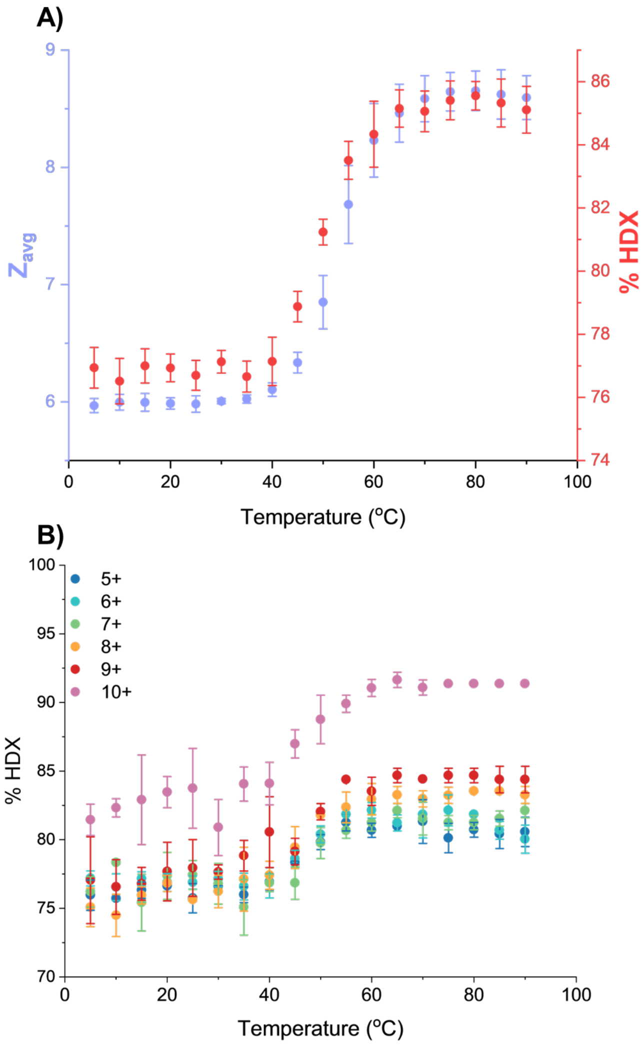


**Figure S8. A)** Comparison of the deuterium uptake of CI-2 as a function of temperature in $D_2O$ with the average charge state and **B)** individual charge states. %HDX was calculated from the weighted average mass from each charge state in the mass spectra for n=3 trials where 100% would indicate that all titratable residues have exchanged. These results indicate that certain titratable residues are hidden at cold temperatures and become exposed as the protein unfolds. For each charge state individually the trend is the same where deuterium uptake by CI-2 increases temperature however the 10+ charge state has a significant amount more uptake indicating largely different structure(s) than the 5-9+ charge states. Since the IM-MS profiles within charge states do not change with temperature the uptake trend provides evidence that the conformations and/or charge states are interconverting at least for the 5-9+ to explain the similar uptake trend

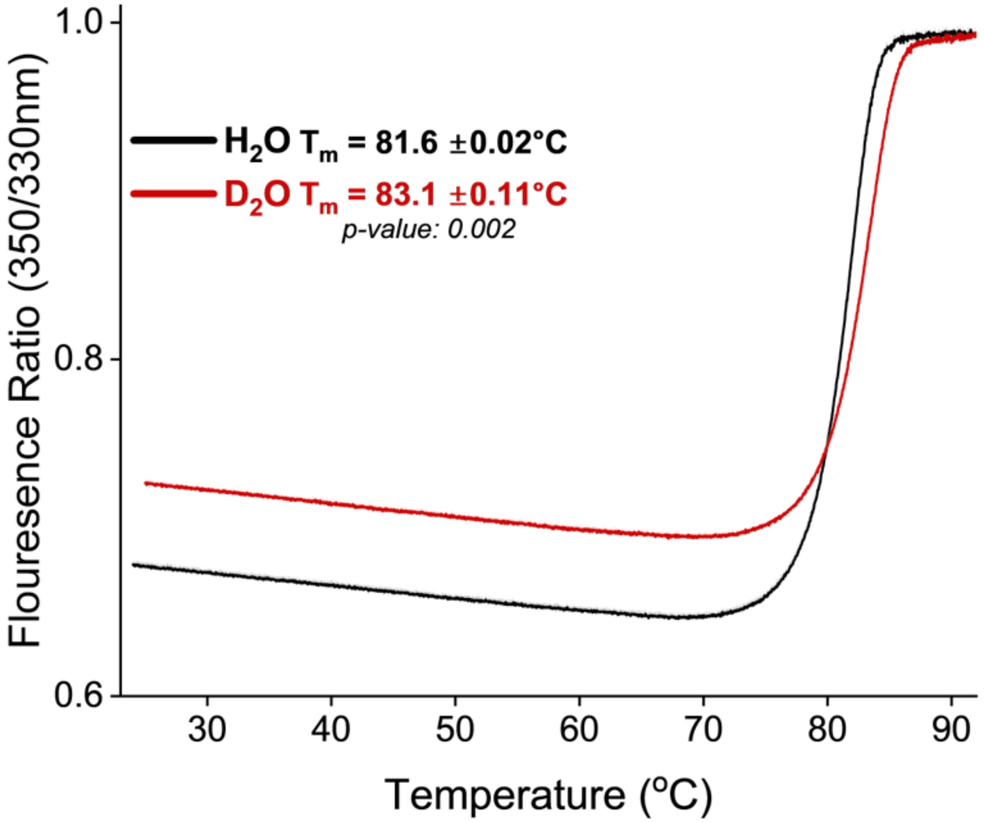


**Figure S9.** Nano DSF results of Bn-B* complex dissociation in both solvents in 10 mM HEPES and 10 mM NaCl at pH/pD 7. The increased melting in $D_2O$ indicates that the results shown in the main text are not an artifact of gas phase MS measurements and represent solution temperature effects. Error bars are present in the plot as faded regions behind the bar of triplicate measurements.

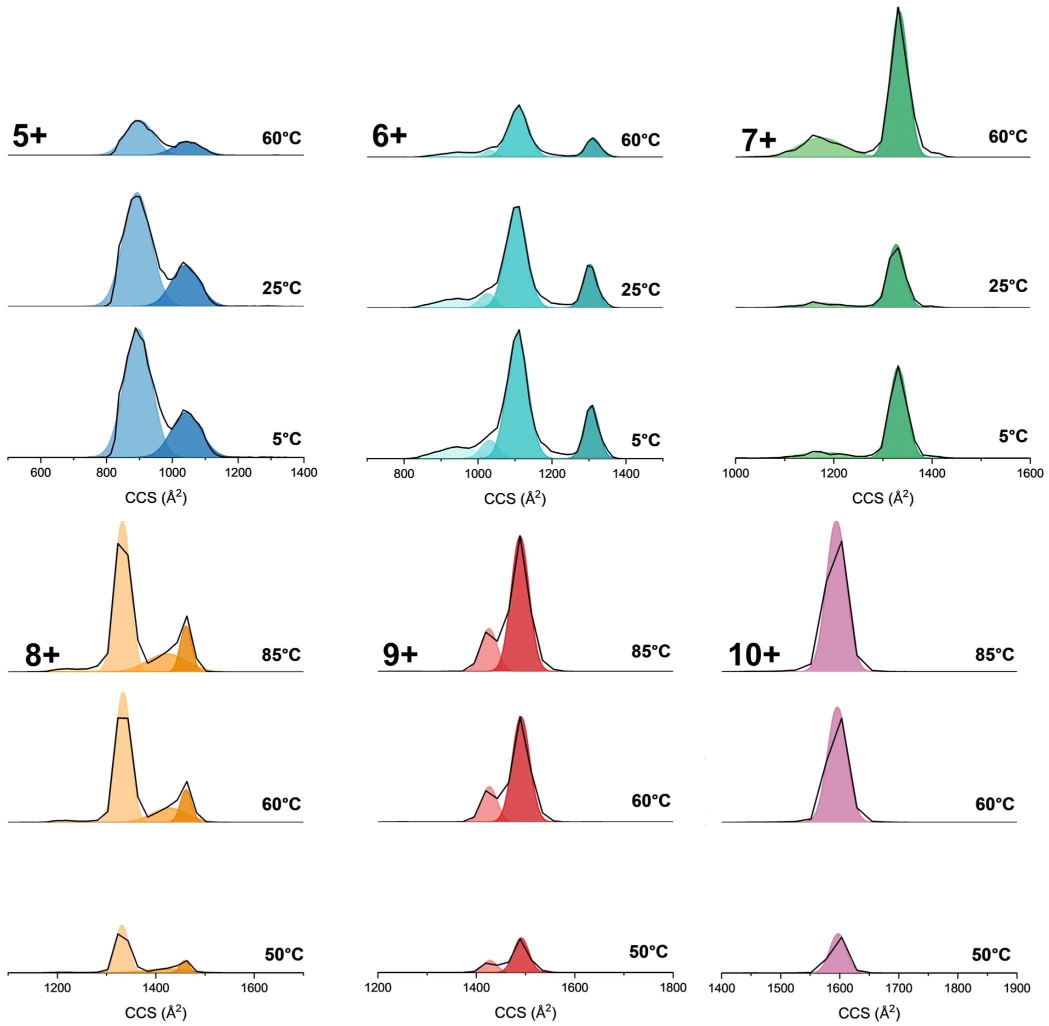


**Figure S10.** $H_2O$ IM-MS data for each CI-2 charge state that shows there is no change of conformation abundance within charge states but as temperature increases the population of the charge state is what changes. Lower charge states (5 & 6+) are inversely proportional to temperature while higher charge state 7-10+ are directly proportional to temperature. The 7+ charge state appears as a potential sensitive intermediate state due to the observable increase in abundance as temperature decreases from 25 ºC to 5 ºC which suggests that there is some slight cold unfolding that occurs. Unfortunately our setup cannot go cold enough to pursue cold unfolding for this system.

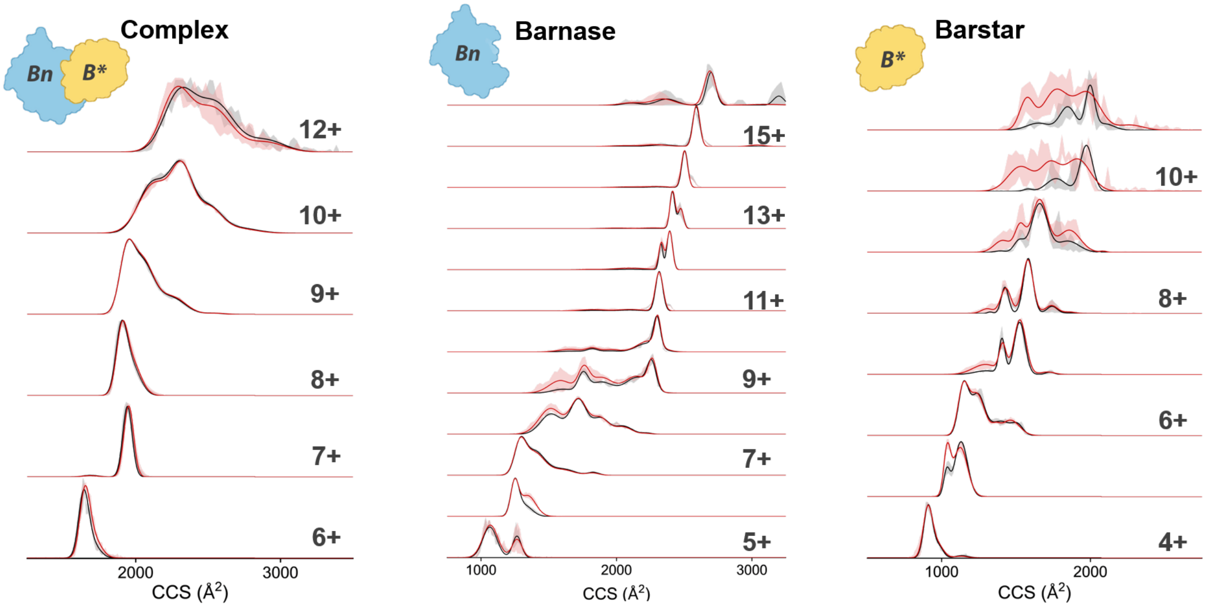


**Figure S11.** Representative raw IM-MS profiles for each Bn-B* complex and individual protein charge state in $H_2O$ (black) and $D_2O$ (red). Faded regions represent standard deviation of replicates (n=3).

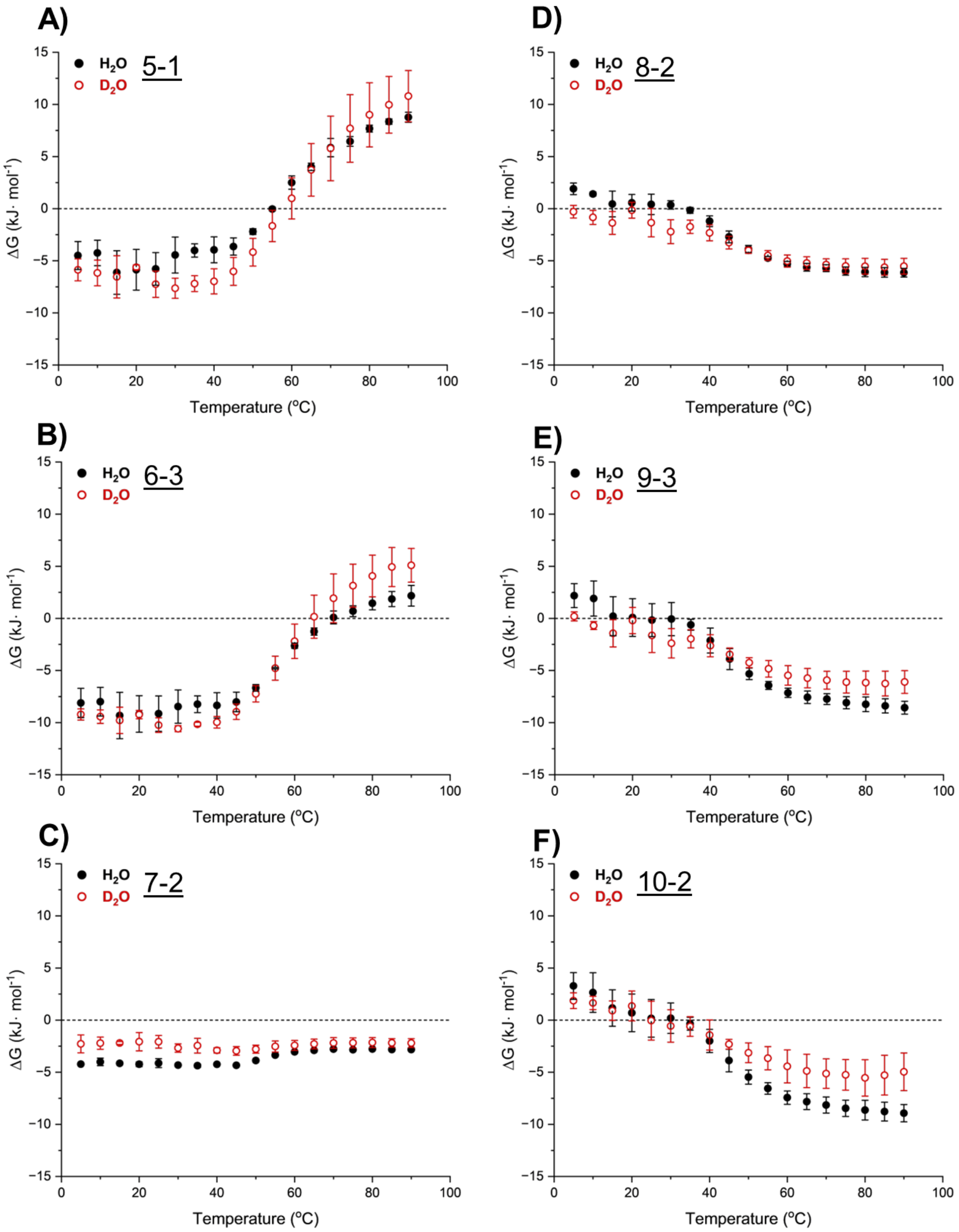


**Figure S12.** Favorability (ΔG) of the most dominant conformation in each CI-2 charge state **A)** 5+ **B)** 6+ **C)** 7+ **D)** 8+ **E)** 9+ **F)** 10+ across all temperatures (5-90 ºC) in both solvents. Notably the higher charge states (9+ & 10+) show less favorability in $D_2O$ at higher temperatures which we attribute to CI-2 in $D_2O$ having a lower SASA and increased abundance of compact conformations that alters the free energy of each state relative to $H_2O$. The decreased favorability in $D_2O$ for the 6+ at higher temperatures indicates there is a higher barrier to unfolding and/or the process may not be as easily reversible.

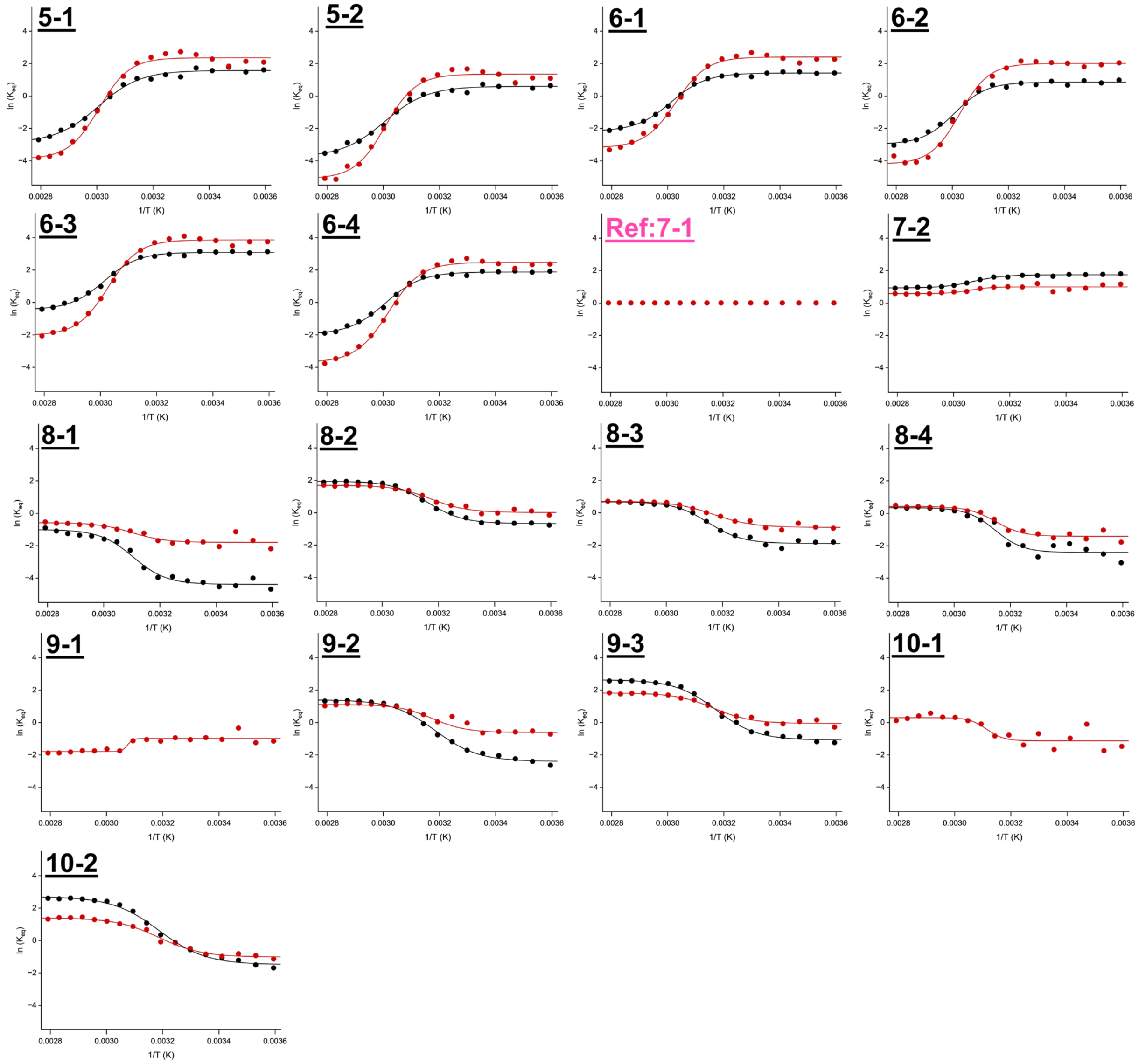


**Figure S13.** Representative Van't Hoff plots (ln($K_{eq}$) vs 1/T (K)) for each CI-2 conformation across temperature for both $H_2O$ (black) and $D_2O$ (red) from a single trial. Fits to the data that were used for all thermodynamic analysis are shown as a solid line. Larger changes in the equilibrium constant are seen for the native conformations (5+ & 6+) in $D_2O$ and for unfolded conformations (8+ - 10+) in $H_2O$.

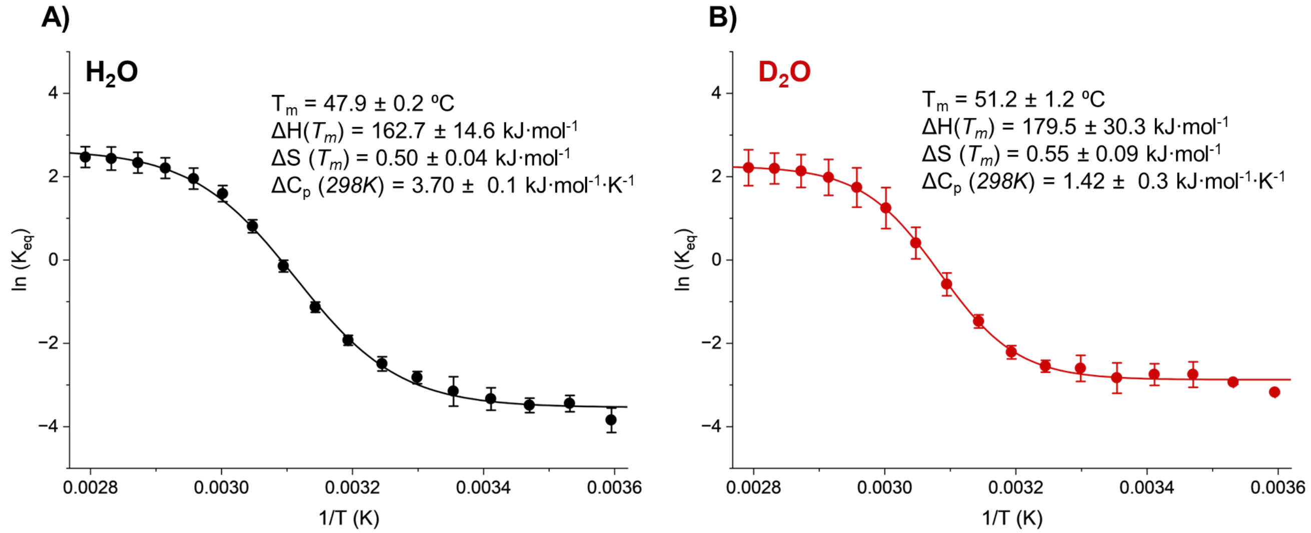


**Figure S14.** Ensemble analysis using the abundance of native (N) conformations (5+ - 7+) vs unfolded (U) conformations (8+ - 10+). The equilibrium constant at each temperature was calculated by $K_{eq} = \frac{\sum U}{\sum N}$ for Van't Hoff analysis in **A)** $H_2O$ and **B)** $D_2O$. Fits to the average data can be seen as dashed lines on both plots. Heat capacity was calculated as the derivate of the enthalpy versus temperature. The results match closely with prior DSC data at pH 2.5 $T_m = 47.1$ ºC $\Delta H(T_m) = 205.9$ kJ·mol$^{-1}$ $\Delta S(T_m) = 0.64$ kJ·mol$^{-1}$ and $\Delta C_p = 3.3$ kJ·mol$^{-1}$·K$^{-1}$.[1] When comparing solvents $D_2O$ has a statistically increased melting temperature (p-value = 0.038) and less positive heat capacity indicating more compact unfolded conformations which aligns well with vT-IM-MS data. Error bars represent standard deviation of replicates (n=3) for all data.

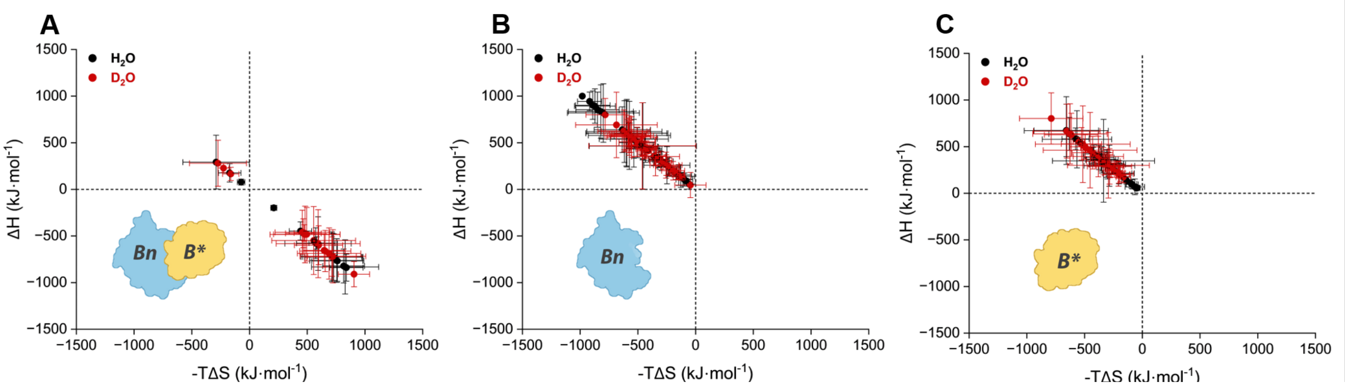


**Figure S15.** Entropy-enthalpy compensation plots (EEC) for each conformation in both solvents for **A)** Bn-B* **B)** Bn and **C)** B*. The individual proteins have opposing favorability trends seen between the solvents which indicates unique interactions with the opposing solvents. Additionally the similar trends of the complex and B* indicate a potentially dominant contribution of the individual protein's conformations to the thermodynamics and thermal stability of the complex.

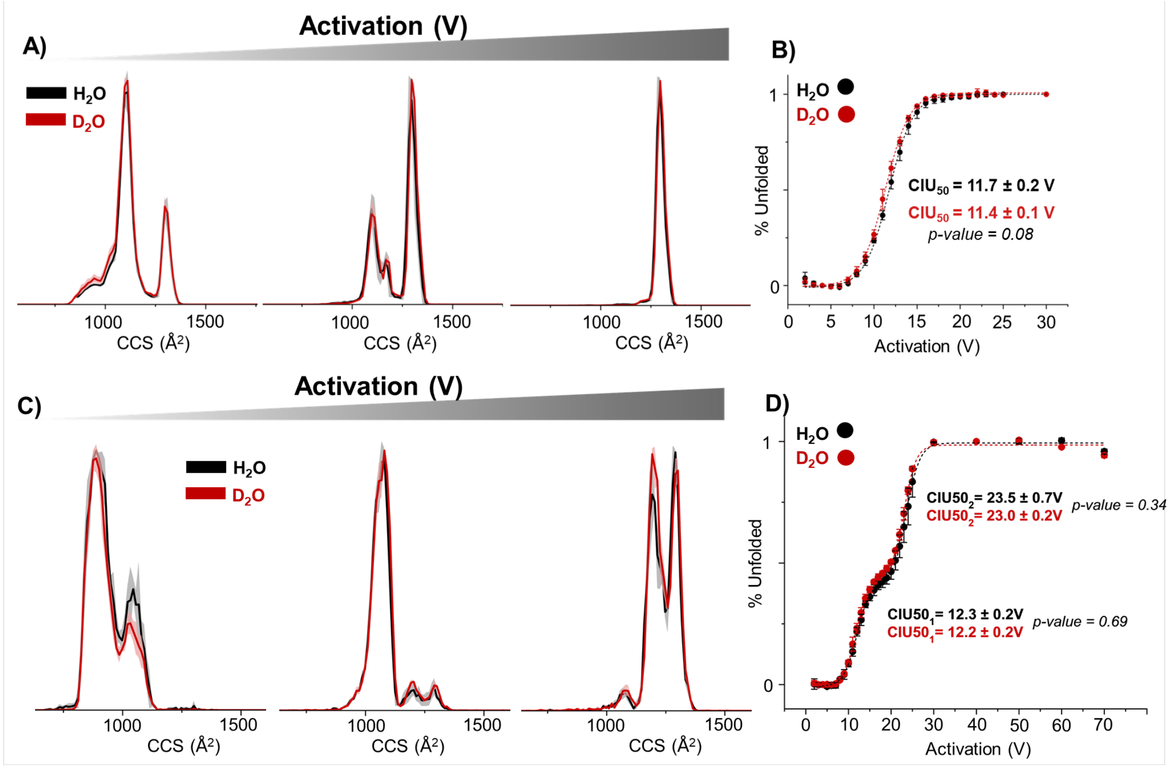


**Figure S16.** IM-MS profiles as a function of increased activation of CI-2 in both $H_2O$ and $D_2O$ for the **A)** 6+ and **C)** 5+ charge state. Calculated change in unfolding of CI-2 **B)** 6+ and **D)** 5+ as a function of activation voltage in 1V increments. There is no statistically significant difference in the $CIU50_1$ or $CIU50_2$ between solvents for both charge states which provides evidence that the deuterium isotopes imparted on CI-2 have little to no effect on the stability of CI-2. In conjunction with the temperature data this data suggests the stabilizing effect seen in $D_2O$ for protein melting are due to altered solvent-protein interactions. Error bars and faded regions represent standard deviation of replicates (n=3).

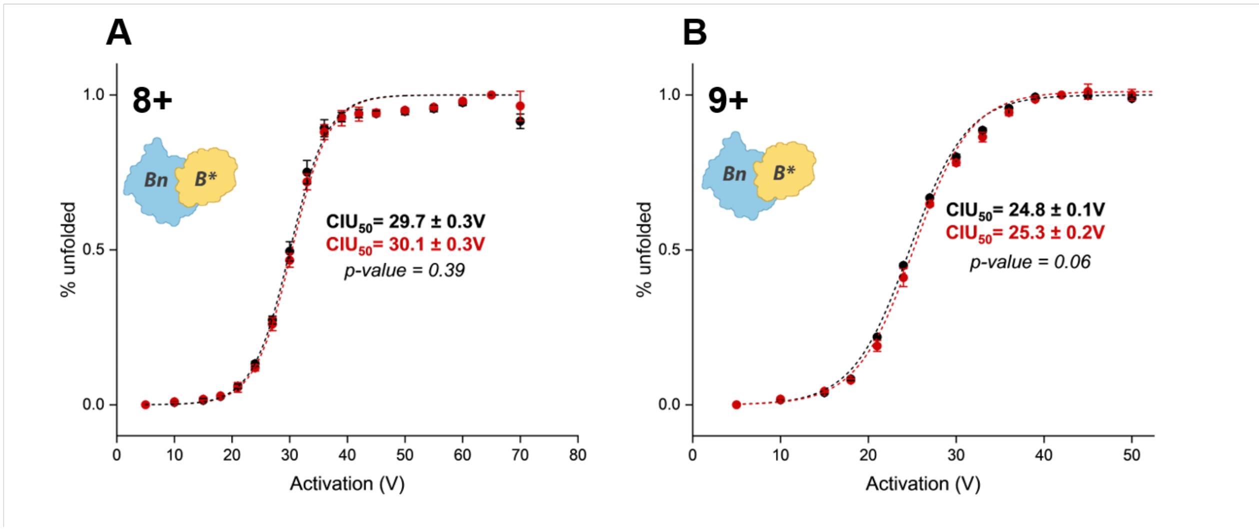


**Figure S17.** CIU profiles as a function of increased activation of Bn-B* in both $H_2O$(black) and $D_2O$(red) for the **A)** 8+ and **C)** 9+ charge state. There is no statistically significant difference in the CIU50 between solvents for both charge states which provides evidence that the deuterium isotopes imparted on Bn-B* have little to no effect on the stability. Error bars represent standard deviation of replicates (n=3).

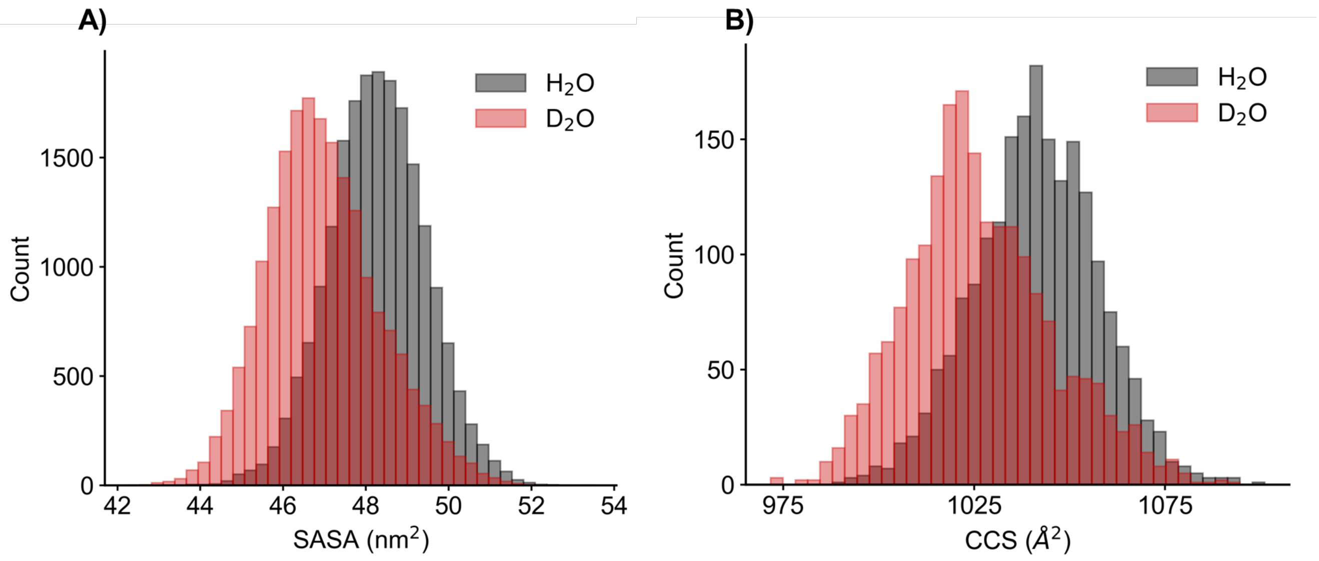


**Figure S18. A)** Solvent accessible surface area (SASA) and **B)** collision cross section (CCS) of CI-2 from the entire trajectory of the MDS in either $H_2O$ or $D_2O$ solvent. These results suggest the protein adopts a more compact conformation in $D_2O$ rather than just a difference in exposed residue SASA. CCS was calculated from each frame of the MDS using IMPACT. [2]

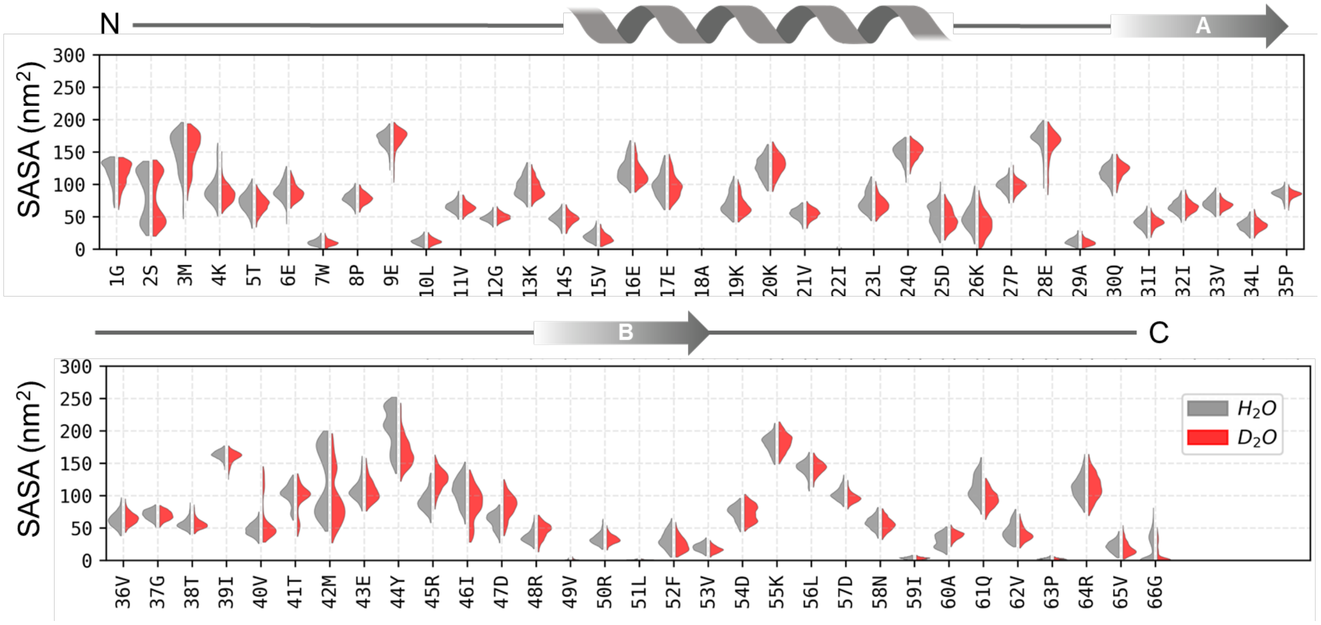


**Figure S19.** Solvent accessible surface area of each CI-2 residue from the entire trajectory of the MDS in either $H_2O$ or $D_2O$ solvent. Secondary structure was taken from the 5 structures closest to the mean of the SASA as seen in panel A of Figure SX. The loop region around the hydrophobic core V40-R48 is the most different between solvents which suggests this region may be important for stability and the difference in conformations accessible. More subtle increased compaction (decreased SASA) and dynamics (smaller FWHM) in $D_2O$ can additionally be seen extending towards the C-terminus (D54-G66) and throughout the loop/ beta sheet A region (D25-I39).

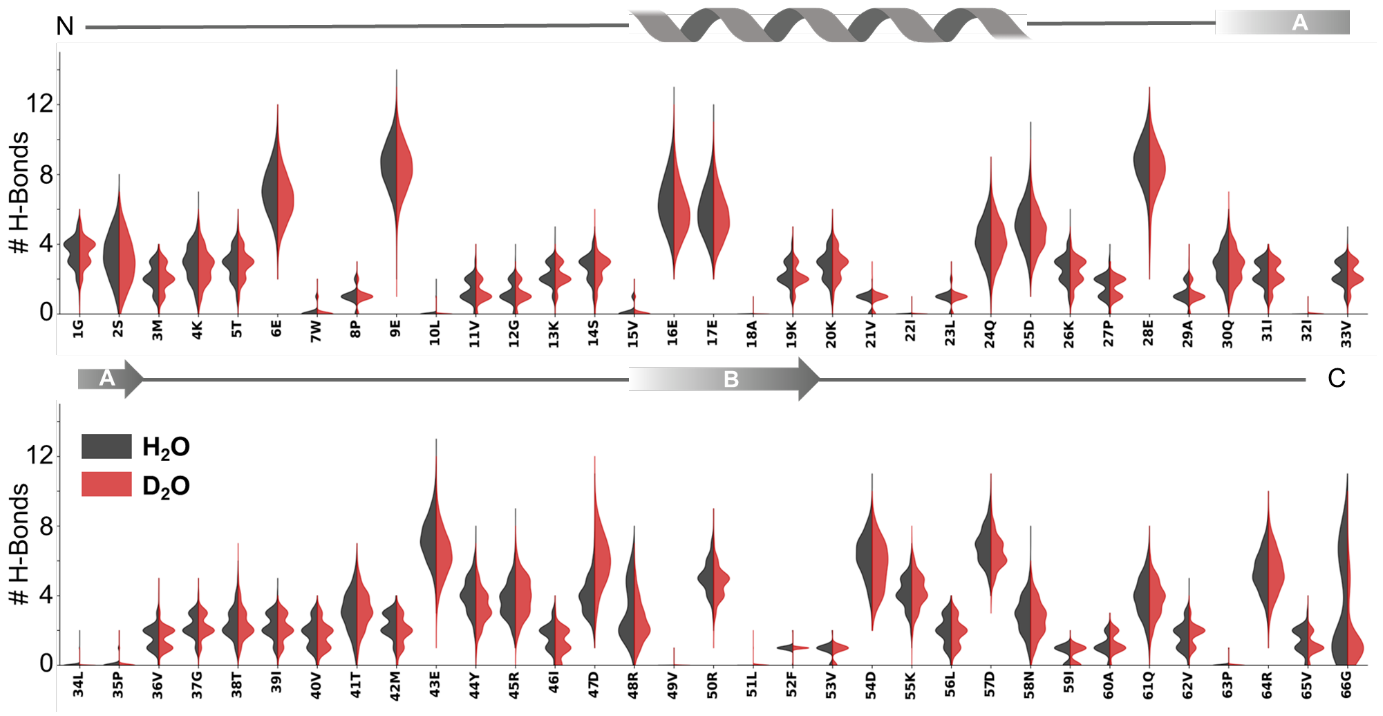


**Figure S20.** Number of solvent-protein hydrogen bonds for each CI-2 residue from the entire trajectory of the MDS in either $H_2O$ or $D_2O$ solvent. Secondary structure was taken from the 5 structures closest to the mean of the SASA as seen in panel A of Figure SX. The results are largely similar to that of SASA residue analysis. The loop region around the hydrophobic core V40-R48 is the most different between solvents which suggests this region is most sensitive to hydration.

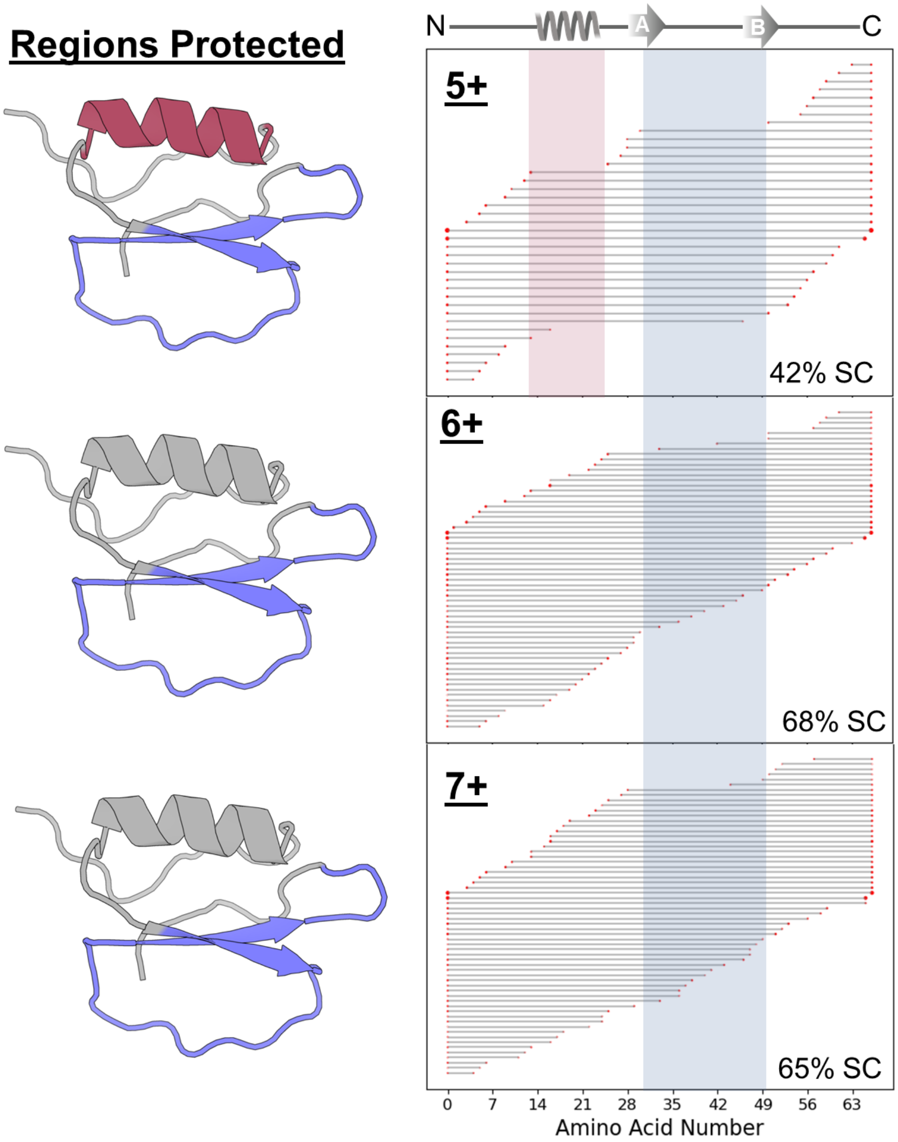


**Figure S21.** Electron capture dissociation data of the 5-7+ charge states of CI-2 and the protected region mapped onto the average MDS structure. Analysis of the c/z ions was completed using ClipsMS (right) and the regions where there are gaps in the sequence coverage is used to indicate protected regions of the proteins structure.[3] The 5+ charge state shows all secondary structures are protected providing evidence they are present at this charge. For the 6+ and 7+ the alpha helix region is no longer protected suggesting it is the first region to unfold as the protein populates higher charge states. Additionally this data supports the beta sheet and loop region 40V-48R is highly important in unfolding and the presence of compact conformations. ECD data was collected on an Agilent 6545XT with an ExD cell.

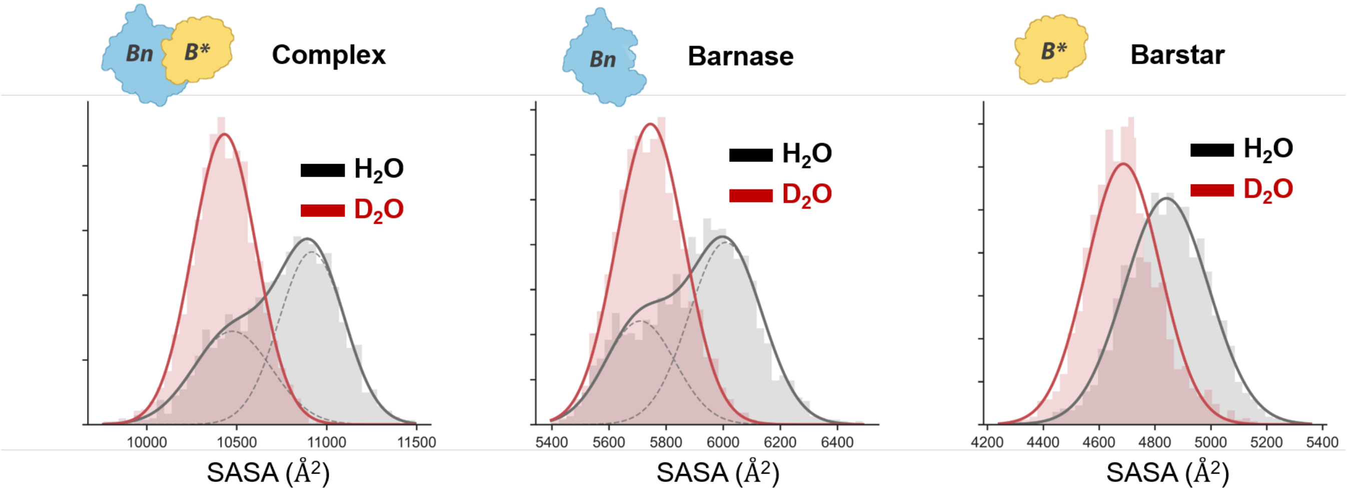


**Figure S22.** Solvent accessible surface area (SASA) of Bn-B* and the individua proteins from the entire trajectory of the MDS in either $H_2O$ or $D_2O$ solvent. These results suggest the protein adopts a more compact conformation in $D_2O$ and is less dynamic due to bimodal distribution found only in $H_2O$.

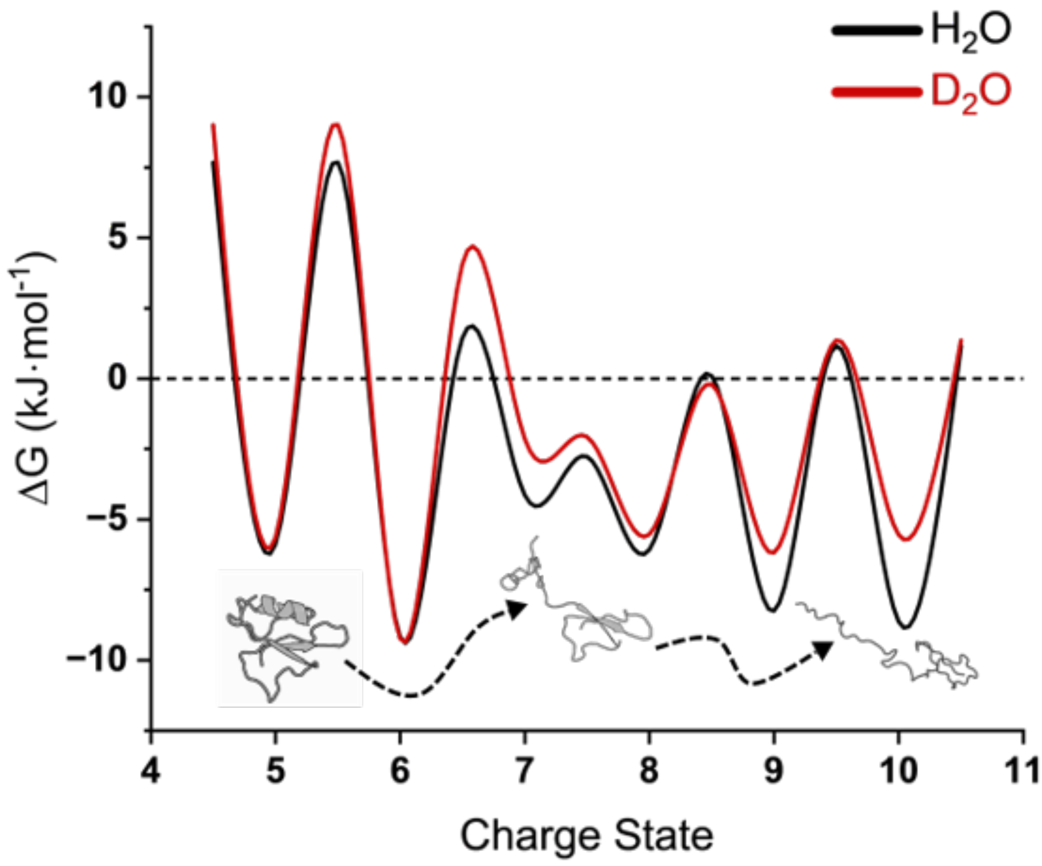


**Figure S23.** Free energy landscape of unfolding for CI-2 in both $H_2O$ (black) and $D_2O$ (red) using the most dominant conformations in each charge state. At low charge states or native conformations the energy wells are similar however the barriers to overcome are higher in $D_2O$ indicating less protein-water interactions. The higher charge states (8-10+) or unfolded conformations have more shallow wells in $D_2O$ which indicate they are less favorable to populate. The differences in landscape between solvents directly indicates the increased Tm in $D_2O$ is due to changes in CI-2 hydration.

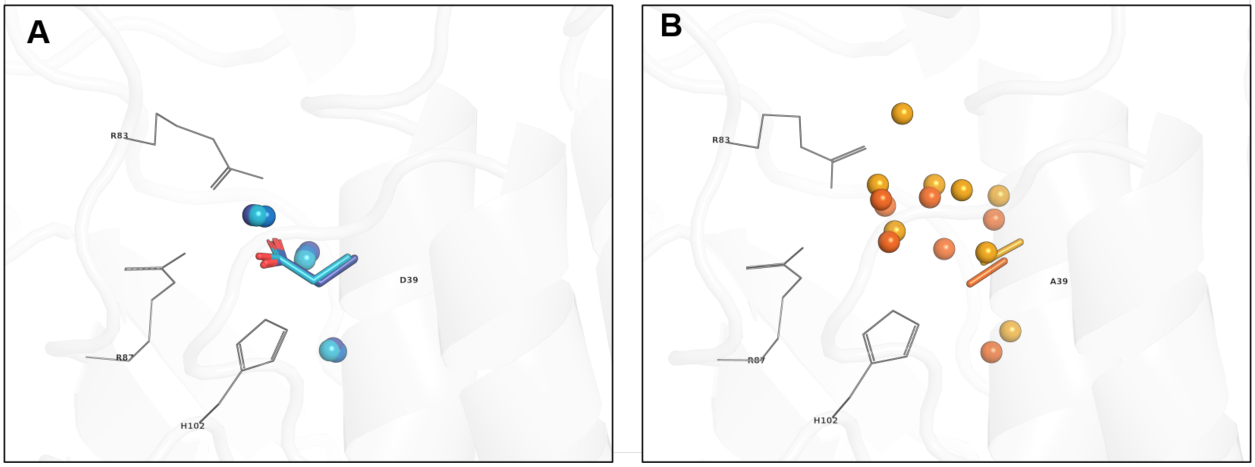


**Figure S24.** Overlayed structures of water interactions with the region that holds Barstar residue **A)** D39 (D40 in our work) pbd: 1X1U and mutant **B)** D39A pbd: 2ZA4. Comparison of the WT and D39A crystal structures suggest that interfacial waters are more consistently positioned in the WT, whereas the mutant exhibits a more heterogeneous distribution of resolved waters, with some water positions not conserved across crystallographically independent chains. The water selection is based on 5 Å of any heavy atom of reference WT residue D39.

**Table S1.** Waters Synapt G2 Parameters for All Experiments

| Synapt G2 Parameters | CI-2 | Bn-B* |
|---|---|---|
| Capillary (kV) | 0.9 – 1.4 | 0.9 – 1.4 |
| Source Temperature (ºC) | 20 | 50 |
| Sampling Cone (V) | 10 | 10 |
| Extraction Cone (V) | 1 | 1 |
| Trap Collision Energy (V) | Off (4) | Off (4) |
| Transfer Collision Energy (V) | Off (0) | Off (0) |
| Trap Gas Flow (mL/min) | 0.40 | 0.40 |
| Helium Cell Gas Flow (mL/min) | 200 | 50 |
| IMS Gas low (mL/min) | 80 | 70 |
| Trap DC Entrance (V) | -4 | 0 |
| Trap DC Bias (V) | 20 | 25 |
| Trap DC (V) | 3 | -2 |
| Trap DC Exit (V) | -3.3 | 3 |
| IMS DC Entrance (V) | 15 | 15 |
| Helium Cell DC (V) | 20.1 | 25 |
| Helium Exit (V) | -5 | -5 |
| IMS Bias (V) | 0 | 0 |
| IMS DC Exit (V) | 0 | 0 |
| Transfer DC Entrance (V) | 4.4 | 2.6 |
| Transfer DC Exit (V) | 9.1 | 15 |
| Source Wave Velocity (m/s) | 200 | Off |
| Source Wave Height (V) | 1 | Off |
| Trap Wave Velocity (m/s) | 400 | Off |
| Trap Wave Height (V) | 4.9 | Off |
| IMS Wave Velocity (m/s) | 400 | 600 |
| IMS Wave Height (V) | 25 | 25 |
| Transfer Wave Velocity (m/s) | 100 | Off |
| Transfer Wave Height (V) | 7 | Off |
| Sample Collection Time (min) | 1.71 (100 scans) | 1.71 (100 scans) |

**Table S2.** Average weighted CCS and FWHM for each CI-2 conformation in both solvents. Error represents standard deviation of replicates (n=3).

| Conformer | $H_2O$ | | $D_2O$ | |
|---|---|---|---|---|
| | CCS | FWHM | CCS | FWHM |
| 5-1 | 895 ± 0.8 Å$^2$ | 98 ± 0.9 Å$^2$ | 894 ± 2.4 Å$^2$ | 92 ± 1.1 Å$^2$ |
| 5-2 | 1040 ± 1.5 Å$^2$ | 105 ± 1.4 Å$^2$ | 1035 ± 6.1 Å$^2$ | 120 ± 6.6 Å$^2$ |
| 6-1 | 947 ± 0.4 Å$^2$ | 126 ± 1.3 Å$^2$ | 944 ± 2.2 Å$^2$ | 126 ± 2.7 Å$^2$ |
| 6-2 | 1030 ± 0.5 Å$^2$ | 50 ± 0.5 Å$^2$ | 1031 ± 0.5 Å$^2$ | 54 ± 1.6 Å$^2$ |
| 6-3 | 1106 ± 0.5 Å$^2$ | 65 ± 1.2 Å$^2$ | 1109 ± 1.6 Å$^2$ | 68 ± 0.4 Å$^2$ |
| 6-4 | 1307 ± 0.1 Å$^2$ | 46 ± 0.2 Å$^2$ | 1310 ± 1.5 Å$^2$ | 47 ± 1.1 Å$^2$ |
| 7-1 (Ref) | 1175 ± 1.2 Å$^2$ | 108 ± 2.7 Å$^2$ | 1177 ± 0.4 Å$^2$ | 116 ± 0.9 Å$^2$ |
| 7-2 | 1331 ± 1.0 Å$^2$ | 41 ± 0.2 Å$^2$ | 1335 ± 0.3 Å$^2$ | 43 ± 0.5 Å$^2$ |
| 8-1 | 1234 ± 3.2 Å$^2$ | 94 ± 12.3 Å$^2$ | 1221 ± 1.8 Å$^2$ | 71 ± 1.8 Å$^2$ |
| 8-2 | 1331 ± 1.5 Å$^2$ | 35 ± 0.8 Å$^2$ | 1335 ± 0.4 Å$^2$ | 38 ± 0.2 Å$^2$ |
| 8-3 | 1425 ± 0.3 Å$^2$ | 86 ± 2.3 Å$^2$ | 1432 ± 1.2 Å$^2$ | 79 ± 0.9 Å$^2$ |
| 8-4 | 1459 ± 1.7 Å$^2$ | 25 ± 1.3 Å$^2$ | 1466 ± 0.7 Å$^2$ | 27 ± 0.2 Å$^2$ |
| 9-1 | | | 1284 ± 19.3 Å$^2$ | 129 ± 35.1 Å$^2$ |
| 9-2 | 1425 ± 2.1 Å$^2$ | 38 ± 1.0 Å$^2$ | 1430 ± 0.6 Å$^2$ | 33 ± 1.5 Å$^2$ |
| 9-3 | 1488 ± 1.7 Å$^2$ | 42 ± 0.6 Å$^2$ | 1495 ± 0.9 Å$^2$ | 55 ± 1.6 Å$^2$ |
| 10-1 | | | 1562 ± 7.4 Å$^2$ | 107 ± 14.3 Å$^2$ |
| 10-2 | 1593 ± 2.1 Å$^2$ | 39 ± 0.7 Å$^2$ | 1600 ± 1.0 Å$^2$ | 38 ± 2.1 Å$^2$ |

**Table S3.** Average melting and formation temperature for each CI-2 conformation in both solvents. Error represents standard deviation of replicates (n=3).

| Conformer | $H_2O$ | $D_2O$ |
|---|---|---|
| 5-1 | 46.7 ± 0.7 ºC | 52.4 ± 1.4 ºC |
| 5-2 | 46.8 ± 0.9 ºC | 52.4 ± 1.7 ºC |
| 6-1 | 50.4 ± 0.3 ºC | 52.1 ± 1.6 ºC |
| 6-2 | 50.1 ± 1.5 ºC | 50.0 ± 1.2 ºC |
| 6-3 | 50.0 ± 0.8 ºC | 51.7 ± 1.5 ºC |
| 6-4 | 49.8 ± 0.7 ºC | 52.1 ± 1.8 ºC |
| 7-1 (Ref) | 29.8 ± 6.2 ºC | 49.4 ± 4.4 ºC |
| 7-2 | 56.4 ± 3.6 ºC | 46.9 ± 2.1 ºC |
| 8-1 | 52.2 ± 2.5 ºC | 54.6 ± 2.1 ºC |
| 8-2 | 48.4 ± 1.3 ºC | 51.7 ± 1.7 ºC |
| 8-3 | 48.4 ± 2.3 ºC | 52.2 ± 2.1 ºC |
| 8-4 | 50.0 ± 2.0 ºC | 52.8 ± 1.3 ºC |
| 9-1 | | 48.0 ± 2.6 ºC |
| 9-2 | 49.6 ± 0.5 ºC | 52.0 ± 1.7 ºC |
| 9-3 | 50.3 ± 0.5 ºC | 52.6 ± 1.6 ºC |
| 10-1 | | 52.3 ± 0.9 ºC |
| 10-2 | 51.2 ± 0.9 ºC | 53.8 ± 1.8 ºC |

**Table S4.** Average weighted CCS and FWHM for each Bn-B* conformation in both solvents. Error represents standard deviation of replicates (n=3).

| Conformer | $H_2O$ | | $D_2O$ | |
|---|---|---|---|---|
| | CCS | FWHM | CCS | FWHM |
| C_6_1 | 1645 ± 5 Å² | 85 ± 6 Å² | 1657 ± 7 Å² | 88 ± 7 Å² |
| C_6_2 | 1713 ± 10 Å² | 107 ± 15 Å² | 1720 ± 10 Å² | 129 ± 11 Å² |
| C_7_1 | 1751 ± 20 Å² | 189 ± 11 Å² | 1775 ± 16 Å² | 197 ± 8 Å² |
| C_7_2 | 1933 ± 9 Å² | 122 ± 26 Å² | 1953 ± 5 Å² | 104 ± 18 Å² |
| C_8_1 | 1909 ± 4 Å² | 92 ± 2 Å² | 1916 ± 4 Å² | 93 ± 2 Å² |
| C_8_2 | 1977 ± 4 Å² | 146 ± 2 Å² | 1985 ± 5 Å² | 149 ± 2 Å² |
| C_9_1 | 1936 ± 2 Å² | 116 ± 2 Å² | 1938 ± 2 Å² | 115 ± 2 Å² |
| C_9_2 | 2043 ± 3 Å² | 187 ± 2 Å² | 2046 ± 4 Å² | 187 ± 2 Å² |
| C_9_3 | 2252 ± 2 Å² | 215 ± 5 Å² | 2256 ± 3 Å² | 218 ± 5 Å² |
| C_9_4 | 2540 ± 6 Å² | 161 ± 7 Å² | 2552 ± 4 Å² | 150 ± 6 Å² |
| C_10_1 | 2115 ± 8 Å² | 226 ± 7 Å² | 2114 ± 6 Å² | 236 ± 6 Å² |
| C_10_2 | 2316 ± 3 Å² | 199 ± 9 Å² | 2320 ± 3 Å² | 193 ± 9 Å² |
| C_10_3 | 2517 ± 6 Å² | 190 ± 7 Å² | 2519 ± 6 Å² | 191 ± 8 Å² |
| C_10_4 | 2709 ± 6 Å² | 296 ± 10 Å² | 2706 ± 5 Å² | 293 ± 11 Å² |
| C_12_1 | 2305 ± 21 Å² | 253 ± 28 Å² | 2311 ± 27 Å² | 253 ± 30 Å² |
| C_12_2 | 2552 ± 25 Å² | 296 ± 9 Å² | 2560 ± 23 Å² | 297 ± 11 Å² |
| C_12_3 | 2908 ± 18 Å² | 293 ± 13 Å² | 2909 ± 14 Å² | 293 ± 11 Å² |

| Conformer | $H_2O$ | | $D_2O$ | |
|---|---|---|---|---|
| | CCS | FWHM | CCS | FWHM |
| Bn_5_1 (Ref) | 1081 ± 14 Å² | 120 ± 5 Å² | 1095 ± 13 Å² | 121 ± 5 Å² |
| Bn_5_2 | 1262 ± 3 Å² | 77 ± 8 Å² | 1259 ± 4 Å² | 90 ± 10 Å² |
| Bn_6_1 | 1248 ± 1 Å² | 71 ± 1 Å² | 1247 ± 1 Å² | 65 ± 1 Å² |
| Bn_6_2 | 1309 ± 4 Å² | 124 ± 7 Å² | 1340 ± 3 Å² | 149 ± 4 Å² |
| Bn_7_1 | 1289 ± 2 Å² | 95 ± 1 Å² | 1290 ± 3 Å² | 95 ± 1 Å² |
| Bn_7_2 | 1393 ± 1 Å² | 179 ± 2 Å² | 1390 ± 2 Å² | 180 ± 2 Å² |
| Bn_7_3 | 1605 ± 2 Å² | 200 ± 1 Å² | 1600 ± 2 Å² | 200 ± 2 Å² |
| Bn_7_4 | 1831 ± 1 Å² | 101 ± 2 Å² | 1828 ± 2 Å² | 113 ± 6 Å² |
| Bn_8_1 | 1517 ± 2 Å² | 171 ± 4 Å² | 1517 ± 6 Å² | 184 ± 5 Å² |
| Bn_8_2 | 1719 ± 2 Å² | 140 ± 1 Å² | 1718 ± 6 Å² | 141 ± 3 Å² |
| Bn_8_3 | 1886 ± 3 Å² | 144 ± 7 Å² | 1880 ± 4 Å² | 150 ± 7 Å² |
| Bn_8_4 | 2050 ± 1 Å² | 151 ± 5 Å² | 2050 ± 1 Å² | 152 ± 5 Å² |
| Bn_8_5 | 2219 ± 5 Å² | 92 ± 8 Å² | 2220 ± 7 Å² | 100 ± 10 Å² |
| Bn_9_1 | 1597 ± 8 Å² | 166 ± 16 Å² | 1591 ± 11 Å² | 185 ± 13 Å² |
| Bn_9_2 | 1755 ± 4 Å² | 79 ± 10 Å² | 1760 ± 6 Å² | 98 ± 15 Å² |
| Bn_9_3 | 1888 ± 12 Å² | 195 ± 13 Å² | 1896 ± 14 Å² | 198 ± 8 Å² |
| Bn_9_4 | 2154 ± 13 Å² | 197 ± 7 Å² | 2147 ± 18 Å² | 195 ± 7 Å² |
| Bn_9_5 | 2258 ± 3 Å² | 76 ± 8 Å² | 2260 ± 5 Å² | 83 ± 9 Å² |
| Bn_10_1 | 1675 ± 14 Å² | 182 ± 17 Å² | 1667 ± 17 Å² | 184 ± 19 Å² |
| Bn_10_2 | 1825 ± 6 Å² | 95 ± 11 Å² | 1824 ± 12 Å² | 105 ± 12 Å² |
| Bn_10_3 | 1950 ± 18 Å² | 124 ± 21 Å² | 1971 ± 15 Å² | 130 ± 17 Å² |
| Bn_10_4 | 2235 ± 4 Å² | 200 ± 2 Å² | 2236 ± 6 Å² | 200 ± 4 Å² |
| Bn_10_5 | 2300 ± 2 Å² | 51 ± 2 Å² | 2304 ± 3 Å² | 53 ± 2 Å² |
| Bn_11_1 | 1859 ± 14 Å² | 153 ± 22 Å² | 1828 ± 26 Å² | 174 ± 19 Å² |
| Bn_11_2 | 2099 ± 6 Å² | 199 ± 5 Å² | 2125 ± 16 Å² | 232 ± 8 Å² |
| Bn_11_3 | 2315 ± 1 Å² | 59 ± 2 Å² | 2317 ± 3 Å² | 63 ± 3 Å² |
| Bn_12_1 | 1832 ± 29 Å² | 96 ± 31 Å² | 1870 ± 22 Å² | 153 ± 23 Å² |
| Bn_12_2 | 2099 ± 5 Å² | 295 ± 17 Å² | 2124 ± 16 Å² | 168 ± 14 Å² |
| Bn_12_3 | 2330 ± 1 Å² | 40 ± 3 Å² | 2334 ± 3 Å² | 46 ± 3 Å² |
| Bn_12_4 | 2394 ± 1 Å² | 46 ± 1 Å² | 2397 ± 3 Å² | 48 ± 2 Å² |
| Bn_13_1 | 1967 ± 26 Å² | 166 ± 32 Å² | 1952 ± 27 Å² | 175 ± 26 Å² |
| Bn_13_2 | 2285 ± 22 Å² | 183 ± 22 Å² | 2270 ± 29 Å² | 196 ± 10 Å² |
| Bn_13_3 | 2414 ± 1 Å² | 44 ± 1 Å² | 2417 ± 2 Å² | 46 ± 2 Å² |
| Bn_13_4 | 2475 ± 2 Å² | 44 ± 2 Å² | 2479 ± 3 Å² | 46 ± 3 Å² |
| Bn_14_1 | 2032 ± 22 Å² | 147 ± 29 Å² | 1996 ± 24 Å² | 165 ± 28 Å² |
| Bn_14_2 | 2278 ± 22 Å² | 189 ± 18 Å² | 2282 ± 16 Å² | 192 ± 12 Å² |
| Bn_14_3 | 2504 ± 1 Å² | 57 ± 4 Å² | 2506 ± 3 Å² | 58 ± 4 Å² |
| Bn_15_1 | 2132 ± 32 Å² | 167 ± 29 Å² | 2062 ± 29 Å² | 168 ± 30 Å² |
| Bn_15_2 | 2343 ± 18 Å² | 165 ± 19 Å² | 2329 ± 24 Å² | 191 ± 14 Å² |
| Bn_15_3 | 2590 ± 2 Å² | 56 ± 3 Å² | 2594 ± 4 Å² | 64 ± 4 Å² |
| Bn_15_4 | 2814 ± 31 Å² | 287 ± 30 Å² | 3020 ± 30 Å² | 141 ± 31 Å² |
| Bn_16_1 | 2146 ± 24 Å² | 144 ± 27 Å² | 2103 ± 17 Å² | 157 ± 27 Å² |
| Bn_16_2 | 2383 ± 15 Å² | 150 ± 25 Å² | 2376 ± 16 Å² | 190 ± 16 Å² |

| Conformer | $H_2O$ | | $D_2O$ | |
|---|---|---|---|---|
| | CCS | FWHM | CCS | FWHM |
| Bn_16_3 | 2691 ± 4 Å² | 68 ± 9 Å² | 2693 ± 7 Å² | 94 ± 10 Å² |
| Bn_16_4 | 3204 ± 5 Å² | 107 ± 14 Å² | 3116 ± 20 Å² | 288 ± 24 Å² |
| BS_4_1 | 905 ± 7 Å² | 70 ± 1 Å² | 903 ± 8 Å² | 72 ± 3 Å² |
| BS_4_2 | 975 ± 7 Å² | 77 ± 7 Å² | 970 ± 8 Å² | 92 ± 10 Å² |
| BS_4_3 | 1152 ± 9 Å² | 85 ± 7 Å² | 1140 ± 10 Å² | 98 ± 4 Å² |
| BS_5_1 | 1034 ± 1 Å² | 47 ± 2 Å² | 1037 ± 1 Å² | 50 ± 1 Å² |
| BS_5_2 | 1134 ± 3 Å² | 110 ± 2 Å² | 1127 ± 3 Å² | 117 ± 3 Å² |
| BS_6_1 | 1154 ± 2 Å² | 80 ± 2 Å² | 1152 ± 2 Å² | 82 ± 2 Å² |
| BS_6_2 | 1252 ± 5 Å² | 91 ± 7 Å² | 1247 ± 3 Å² | 104 ± 6 Å² |
| BS_6_3 | 1393 ± 5 Å² | 155 ± 15 Å² | 1373 ± 6 Å² | 102 ± 15 Å² |
| BS_6_4 | 1505 ± 6 Å² | 102 ± 9 Å² | 1474 ± 6 Å² | 131 ± 7 Å² |
| BS_7_1 | 1298 ± 5 Å² | 166 ± 16 Å² | 1294 ± 9 Å² | 186 ± 11 Å² |
| BS_7_2 | 1408 ± 3 Å² | 49 ± 7 Å² | 1409 ± 10 Å² | 61 ± 17 Å² |
| BS_7_3 | 1525 ± 1 Å² | 81 ± 4 Å² | 1527 ± 4 Å² | 97 ± 5 Å² |
| BS_7_4 | 1727 ± 6 Å² | 64 ± 13 Å² | 1714 ± 5 Å² | 109 ± 9 Å² |
| BS_8_1 | 1306 ± 15 Å² | 52 ± 10 Å² | 1295 ± 11 Å² | 90 ± 19 Å² |
| BS_8_2 | 1427 ± 3 Å² | 64 ± 8 Å² | 1430 ± 5 Å² | 83 ± 11 Å² |
| BS_8_3 | 1581 ± 2 Å² | 85 ± 7 Å² | 1581 ± 3 Å² | 80 ± 9 Å² |
| BS_8_4 | 1744 ± 4 Å² | 69 ± 8 Å² | 1732 ± 7 Å² | 107 ± 10 Å² |
| BS_8_5 | 1830 ± 19 Å² | 74 ± 21 Å² | 1848 ± 11 Å² | 101 ± 17 Å² |
| BS_9_1 | 1387 ± 14 Å² | 85 ± 6 Å² | 1399 ± 20 Å² | 127 ± 19 Å² |
| BS_9_2 | 1517 ± 7 Å² | 81 ± 5 Å² | 1509 ± 18 Å² | 85 ± 8 Å² |
| BS_9_3 | 1666 ± 5 Å² | 140 ± 16 Å² | 1663 ± 7 Å² | 150 ± 15 Å² |
| BS_9_4 | 1852 ± 6 Å² | 159 ± 16 Å² | 1885 ± 9 Å² | 121 ± 19 Å² |
| BS_9_5 | 1977 ± 29 Å² | 123 ± 31 Å² | 2021 ± 42 Å² | 99 ± 29 Å² |
| BS_10_1 | 1569 ± 18 Å² | 100 ± 24 Å² | 1536 ± 19 Å² | 150 ± 32 Å² |
| BS_10_2 | 1771 ± 8 Å² | 127 ± 18 Å² | 1754 ± 22 Å² | 141 ± 11 Å² |
| BS_10_3 | 1974 ± 6 Å² | 94 ± 13 Å² | 1945 ± 14 Å² | 138 ± 24 Å² |
| BS_11_1 | 1664 ± 25 Å² | 169 ± 21 Å² | 1589 ± 18 Å² | 117 ± 23 Å² |
| BS_11_2 | 1870 ± 14 Å² | 152 ± 23 Å² | 1814 ± 29 Å² | 174 ± 23 Å² |
| BS_11_3 | 2008 ± 15 Å² | 82 ± 24 Å² | 1996 ± 22 Å² | 137 ± 33 Å² |
| BS_11_4 | 2143 ± 45 Å² | 144 ± 23 Å² | 2187 ± 54 Å² | 152 ± 27 Å² |

**Table S5.** Average melting and formation temperature for each Bn-B* conformation in both solvents. Error represents standard deviation of replicates (n=3).

| Conformer | $H_2O$ | $D_2O$ |
|---|---|---|
| C_6_1 | 71.4 ± 0.4 °C | 75.8 ± 0.8 °C |
| C_6_2 | 66.4 ± 2.4 °C | 74.7 ± 0.8 °C |
| C_7_1 | 71.6 ± 0.5 °C | 75.1 ± 0.1 °C |
| C_7_2 | 72.1 ± 0.8 °C | 78.7 ± 5.4 °C |
| C_8_1 | 73.2 ± 1.4 °C | 79.7 ± 4.6 °C |
| C_8_2 | 73.7 ± 0.8 °C | 79.5 ± 4.8 °C |

| Conformer | $H_2O$ | $D_2O$ |
|---|---|---|
| C_9_1 | 74.4 ± 1.0 °C | 79.4 ± 3.3 °C |
| C_9_2 | 74.4 ± 1.1 °C | 79.1 ± 3.5 °C |
| C_9_3 | 74.6 ± 1.1 °C | 79.0 ± 3.5 °C |
| C_9_4 | 74.8 ± 0.7 °C | 80.2 ± 2.9 °C |
| C_10_1 | 74.0 ± 1.4 °C | 78.8 ± 3.7 °C |
| C_10_2 | 75.1 ± 0.8 °C | 79.2 ± 3.1 °C |
| C_10_3 | 74.2 ± 1.5 °C | 79.3 ± 3.4 °C |
| C_10_4 | 75.3 ± 1.0 °C | 79.6 ± 3.0 °C |
| C_12_1 | 64.9 ± 0.7 °C | 73.1 ± 4.7 °C |
| C_12_2 | 71.9 ± 5.1 °C | 73.6 ± 1.1 °C |
| C_12_3 | 76.3 ± 0.8 °C | 73.2 ± 2.1 °C |
| Bn_5_1 (Ref) | 73.4 ± 2.1 °C | 74.6 ± 0.4 °C |
| Bn_5_2 | 62.6 ± 1.3 °C | 73.4 ± 4.3 °C |
| Bn_6_1 | 63.6 ± 2.1 °C | 69.7 ± 0.3 °C |
| Bn_6_2 | 64.4 ± 1.1 °C | 68.7 ± 0.2 °C |
| Bn_7_1 | 63.5 ± 1.2 °C | 67.9 ± 1.6 °C |
| Bn_7_2 | 62.7 ± 2.5 °C | 68.8 ± 0.4 °C |
| Bn_7_3 | 63.0 ± 2.6 °C | 68.9 ± 0.7 °C |
| Bn_7_4 | 63.1 ± 2.7 °C | 68.7 ± 0.8 °C |
| Bn_8_1 | 63.5 ± 2.2 °C | 70.4 ± 1.1 °C |
| Bn_8_2 | 63.2 ± 2.5 °C | 70.1 ± 0.9 °C |
| Bn_8_3 | 63.4 ± 2.4 °C | 70.0 ± 0.4 °C |
| Bn_8_4 | 63.8 ± 1.7 °C | 69.8 ± 0.6 °C |
| Bn_8_5 | 63.4 ± 3.7 °C | 71.1 ± 0.9 °C |
| Bn_9_1 | 64.6 ± 2.1 °C | 73.7 ± 3.6 °C |
| Bn_9_2 | 61.9 ± 3.3 °C | 70.5 ± 1.9 °C |
| Bn_9_3 | 63.8 ± 0.7 °C | 69.1 ± 3.9 °C |
| Bn_9_4 | 63.7 ± 3.2 °C | 69.2 ± 0.3 °C |
| Bn_9_5 | 61.4 ± 1.3 °C | 67.6 ± 1.8 °C |
| Bn_10_1 | 66.0 ± 0.2 °C | 75.5 ± 8.3 °C |
| Bn_10_2 | 63.8 ± 3.1 °C | 71.4 ± 1.2 °C |
| Bn_10_3 | 66.1 ± 2.5 °C | 70.2 ± 4.2 °C |
| Bn_10_4 | 63.1 ± 2.7 °C | 70.0 ± 0.9 °C |
| Bn_10_5 | 63.3 ± 1.2 °C | 69.0 ± 1.1 °C |
| Bn_11_1 | 67.3 ± 0.5 °C | 75.0 ± 3.1 °C |
| Bn_11_2 | 64.3 ± 1.5 °C | 70.8 ± 1.2 °C |
| Bn_11_3 | 63.6 ± 0.7 °C | 68.8 ± 1.3 °C |
| Bn_12_1 | 83.5 ± 2.1 °C | 75.4 ± 4.4 °C |
| Bn_12_2 | 65.0 ± 3.3 °C | 73.0 ± 1.8 °C |
| Bn_12_3 | 63.8 ± 1.1 °C | 70.0 ± 0.2 °C |
| Bn_12_4 | 61.5 ± 2.6 °C | 68.0 ± 1.2 °C |
| Bn_13_1 | 85.0 ± 0.1 °C | 75.4 ± 4.8 °C |
| Bn_13_2 | 60.6 ± 0.9 °C | 74.1 ± 2.8 °C |
| Bn_13_3 | 61.5 ± 2.4 °C | 69.6 ± 1.8 °C |

| Conformer | $H_2O$ | $D_2O$ |
|---|---|---|
| Bn_13_4 | 62.7 ± 2.4 °C | 68.9 ± 1.1 °C |
| Bn_14_1 | 67.7 ± 2.9 °C | 76.3 ± 7.7 °C |
| Bn_14_2 | 63.3 ± 1.0 °C | 76.8 ± 3.8 °C |
| Bn_14_3 | 61.8 ± 3.0 °C | 67.3 ± 3.6 °C |
| Bn_15_1 | 60.9 ± 1.4 °C | 77.3 ± 5.4 °C |
| Bn_15_2 | 71.2 ± 11.8 °C | 78.6 ± 5.7 °C |
| Bn_15_3 | 61.1 ± 1.0 °C | 66.3 ± 5.5 °C |
| Bn_15_4 | 63.8 ± 3.4 °C | 72.4 ± 1.5 °C |
| Bn_16_1 | 63.6 ± 2.8 °C | 75.7 ± 8.4 °C |
| Bn_16_2 | 62.5 ± 2.8 °C | 77.4 ± 6.8 °C |
| Bn_16_3 | 66.4 ± 2.6 °C | 68.2 ± 1.3 °C |
| Bn_16_4 | 73.6 ± 3.2 °C | 71.9 ± 0.9 °C |
| BS_4_1 | 72.4 ± 1.2 °C | 73.0 ± 1.0 °C |
| BS_4_2 | 70.1 ± 0.1 °C | 73.9 ± 1.3 °C |
| BS_4_3 | 60.0 ± 0.1 °C | 70.8 ± 2.7 °C |
| BS_5_1 | 64.2 ± 2.1 °C | 69.6 ± 0.3 °C |
| BS_5_2 | 62.7 ± 0.9 °C | 68.3 ± 0.6 °C |
| BS_6_1 | 61.0 ± 1.2 °C | 67.4 ± 1.4 °C |
| BS_6_2 | 60.9 ± 0.8 °C | 68.2 ± 0.9 °C |
| BS_6_3 | 63.5 ± 1.4 °C | 68.7 ± 1.3 °C |
| BS_6_4 | 61.3 ± 1.6 °C | 68.5 ± 0.7 °C |
| BS_7_1 | 62.6 ± 1.8 °C | 72.5 ± 3.5 °C |
| BS_7_2 | 63.6 ± 3.1 °C | 71.6 ± 1.7 °C |
| BS_7_3 | 64.8 ± 2.0 °C | 70.1 ± 1.2 °C |
| BS_7_4 | 68.1 ± 4.4 °C | 72.0 ± 0.6 °C |
| BS_8_1 | 60.4 ± 0.8 °C | 75.4 ± 7.0 °C |
| BS_8_2 | 64.0 ± 3.5 °C | 71.2 ± 1.1 °C |
| BS_8_3 | 61.4 ± 1.3 °C | 71.3 ± 2.2 °C |
| BS_8_4 | 67.0 ± 6.1 °C | 67.0 ± 4.2 °C |
| BS_8_5 | 68.3 ± 0.5 °C | 72.9 ± 2.3 °C |
| BS_9_1 | 69.7 ± 6.1 °C | 71.5 ± 4.3 °C |
| BS_9_2 | 67.8 ± 8.5 °C | 76.2 ± 1.1 °C |
| BS_9_3 | 65.1 ± 0.8 °C | 70.9 ± 2.4 °C |
| BS_9_4 | 67.0 ± 3.7 °C | 71.7 ± 3.4 °C |
| BS_9_5 | 63.3 ± 3.2 °C | 71.3 ± 2.1 °C |
| BS_10_1 | 84.9 ± 0.1 °C | 77.8 ± 2.0 °C |
| BS_10_2 | 64.4 ± 4.0 °C | 76.4 ± 6.9 °C |
| BS_10_3 | 63.0 ± 2.9 °C | 71.2 ± 1.8 °C |
| BS_11_1 | 85.0 ± 0.1 °C | 77.7 ± 6.2 °C |
| BS_11_2 | 67.1 ± 2.9 °C | 75.1 ± 2.0 °C |
| BS_11_3 | 62.4 ± 4.1 °C | 71.6 ± 1.3 °C |
| BS_11_4 | 65.9 ± 5.1 °C | 72.1 ± 2.9 °C |

(1) Jackson S. E.; Fersht A. R. Folding of chymotrypsin inhibitor 2. 1. Evidence for a two-state transition. *Biochemistry* **1991** *30* (43) 10428–10435. DOI: 10.1021/bi00107a010.
(2) Erik; Matteo; Carol; Andrew; Justin. Collision Cross Sections for Structural Proteomics. *Structure* **2015** *23* (4) 791–799. DOI: 10.1016/j.str.2015.02.010.
(3) Lantz C.; Zenaidee M. A.; Wei B.; Hemminger Z.; Ogorzalek Loo R. R.; Loo J. A. ClipsMS: An Algorithm for Analyzing Internal Fragments Resulting from Top-Down Mass Spectrometry. *Journal of Proteome Research* **2021** *20* (4) 1928–1935. DOI: 10.1021/acs.jproteome.0c00952.
